# Type-IV pili tune an adhesion-migration trade-off during surface colonization of *Pseudomonas aeruginosa*

**DOI:** 10.1101/2023.05.09.538458

**Authors:** Ahmet Nihat Simsek, Matthias D. Koch, Joseph E. Sanfilippo, Zemer Gitai, Gerhard Gompper, Benedikt Sabass

**Affiliations:** Theoretical Physics of Living Matter, Institute of Biological Information Processes, Forschungszentrum Jülich, 52425 Jülich, Germany; Institute for Infectious Diseases and Zoonoses, Department of Veterinary Sciences, Ludwig-Maximilians-Universität München, 80752 Munich, Germany; Department of Biology, Texas A&M University, College Station, Texas 77843, USA; Lewis-Sigler Institute for Integrative Genomics, Princeton University, NJ 08544, USA; Department of Molecular Biology, Princeton University, NJ 08544, USA; Department of Biochemistry, University of Illinois at Urbana-Champaign, Urbana, IL 61801, USA

## Abstract

Bacterial pathogenicity relies on both firm surface adhesion and cell dissemination. How twitching bacteria resolve the fundamental contradiction between adhesion and migration is unknown. To address this question, we employ live-cell imaging of type-IV pili (T4P) and therewith construct a comprehensive mathematical model of *Pseudomonas aeruginosa* migration. The data show that only 10% to 50% of T4P bind to substrates and contribute to migration through random extension and retraction. Individual T4P do not display a measurable sensory response to surfaces, but their number increases on cellular surface contact. Attachment to surfaces is mediated, besides T4P, by passive adhesive forces acting on the cell body. Passive adhesions slow down cell migration and result in local random motion on short time scales, which is followed by directionally persistent, superdiffusive motion on longer time scales. Moreover, passive adhesions strongly enhance surface attachment under shear flow. Δ*pilA* mutants, which produce no T4P, robustly stick to surfaces under shear flow. In contrast, rapidly migrating Δ*pilH* cells, which produce an excessive number of T4P, are easily detached by shear. Wild-type cells sacrifice migration speed for robust surface attachment by maintaining a low number of active pili. The different cell strains pertain to disjunct regimes in a generic adhesion-migration trait space. Depending on the nature of the adhesion structures, adhesion and migration are either compatible or a trade-off is required for efficient bacterial surface colonization under different conditions.

---

Most biological processes involving cell migration fundamentally depend on how cells balance static adhesion and migration. Examples include the epithelial mesenchymal transition, cancer metastasis [1, 2], immune-cell migration [3], and bacterial infection [4–7]. In the case of bacteria, strong adhesion to surfaces is required for the formation of mechanically robust biofilms. However, surface migration may also be required to explore the environment and to produce dynamic biofilm structures. How can adhesion and migration be optimized simultaneously? We study this generic problem by focusing on the paradigmatic surface migration of *Pseudomonas aeruginosa*, see Fig. 1a.

**Figure 1.**
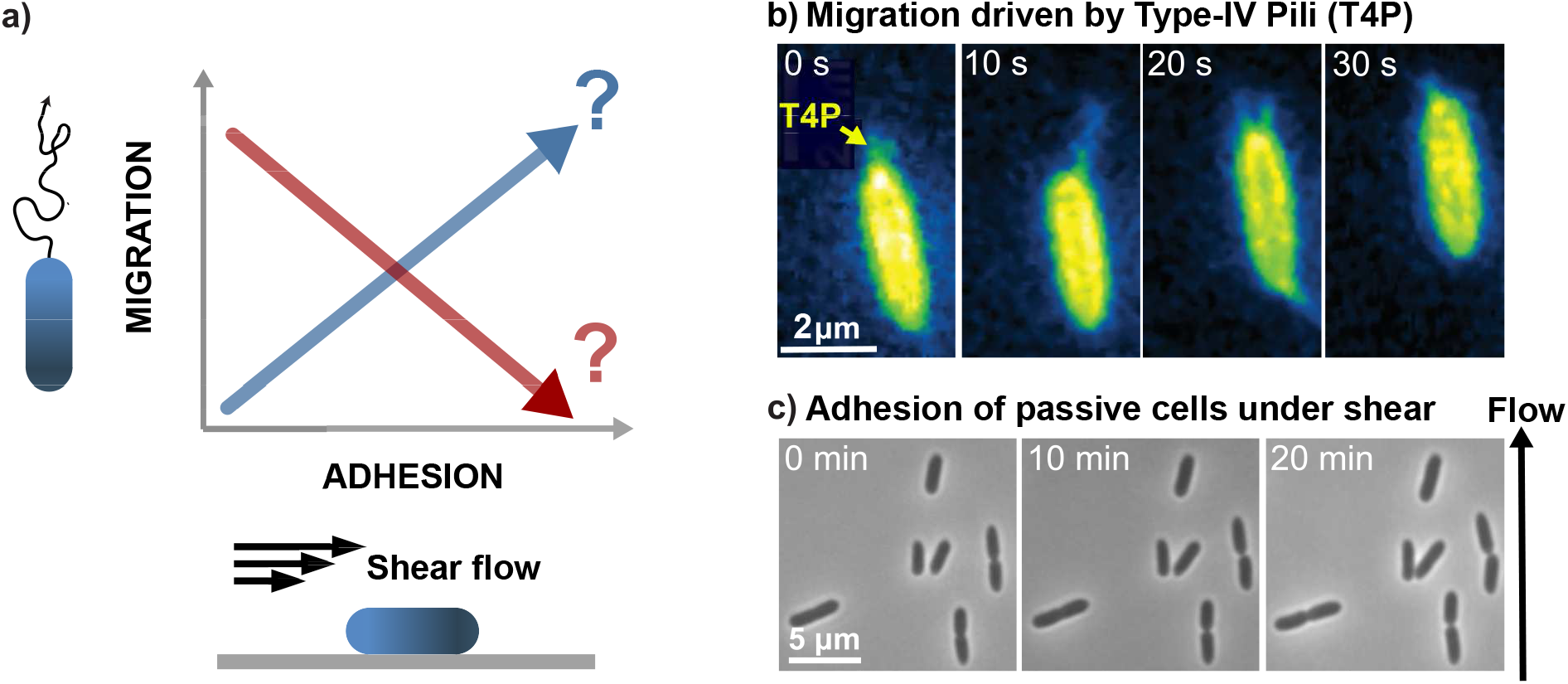
a) Migrating cells under mechanical shear have to balance attachment and detachment to remain at the surface. Can cells simultaneously optimize migration and surface adhesion? b) Live-cell imaging of labeled type IVa pili (T4P) of *P. aeruginosa* for quantification of extension-retraction dynamics during migration. c) Δ*pilA* mutants devoid of T4P stick to the surface under shear flow. Fluid shear stress is 1 Pa.

*P. aeruginosa* is an opportunistic pathogen and the causative agent of various nosocomial infections. Its prominent role as a pathogen is predicated on its ability to colonize a wide variety of environments, including medical catheters, food, and host tissues. *P. aeruginosa* employs dynamic extracellular filaments called type-IVa pili (T4P) to migrate on these surfaces [8–12], see Fig. 1b. T4P-driven motion is called twitching. The filaments are anchored to a molecular machinery in the cell envelope [13], where the ATPase PilB drives pilus polymerzation while pilus retraction is driven by the ATPases PilT and PilU. When pili attach to the extracellular environment, retraction generates a pull with measured stall forces around 30 pN and maximum forces around 150 pN [14]. Extracellularly, *P. aeruginosa* T4P attach to most anorganic or organic materials, including glass and hydrogels, with reported bond strengths up to 95 pN [15]. Thus, T4P-driven migration requires sequential attachment, retraction and detachment of filaments. While the dynamics of freely extending and retracting T4P is well-characterized [16], the stochastic interaction of T4P with surfaces during the twitching process is hardly understood.

*Pseudomonas* infections are often found in environments where shear forces are present, such as the urinary tract. How *P. aeruginosa* adheres to a surface under such conditions remains unclear as well. Using microfluidic setups, it has been shown that shear flow can align cells with the T4P pointing upstream for some bacterial strains. Alignment of T4P leads to upstream migration [17]. However, we find that *P. aeruginosa* mutants without T4P also stick to surfaces under flow, see Fig. 1c. *P. aeruginosa* can adhere passively to a surface through physiochemical forces, through fimbriae expressed in the *cupA* gene cluster[18, 19], and through exopolysaccharides, including those encoded in the *pel* /*psl* gene clusters [20, 21]. Although such passive adhesive forces may play a central role for surface colonization, little is known about their role for adhesion and twitching.

To find out whether bacterial micromechanics are optimised more for migration or for adhesion, we employ fluorescent labeling of T4P in the PAO1 strain [16, 22] and quantify the balance of attachment, detachment, and mechanical activity under different conditions. With this new data, we achieve a comprehensive mathematical model that links whole-cell migration to the dynamics of molecular T4P motors. A central discovery is the existence of a complex adhesion-migration trait space where *P. aeruginosa* realizes a trade-off by maintaining only low numbers of T4P in combination with passive adhesions.

## I. RESULTS

### A. Multiscale model

To derive a model combining the detailed stochastic dynamics of individual T4P with a cell-wide force balance, we analyze fluorescently labeled T4P for cells on different substrates.

#### Quantification of T4P activity on surfaces

T4P stochastically switch between extension, idle, and retraction states, which is explained by the competitive binding of PilB and PilT to the transmembrane complex [16]. A diagram of T4P states is shown in Fig. 2a. For adherent cells, the diagram must be extended to include T4P-surface interaction. For this purpose, we compare T4P statistics on substrates with results for bacteria that are held with an optical trap in liquid suspension, see Fig. 2b. Figure 2c demonstrates that the pilus lengths obey single-exponential distributions for all conditions. For cells on surfaces, T4P are shorter than for cells that are held in liquid suspension with mean lengths of 0.45 *μ*m and 0.77 *μ*m, respectively, see Fig. 2c inset and Supplementary Fig. S4.

**Figure 2.**
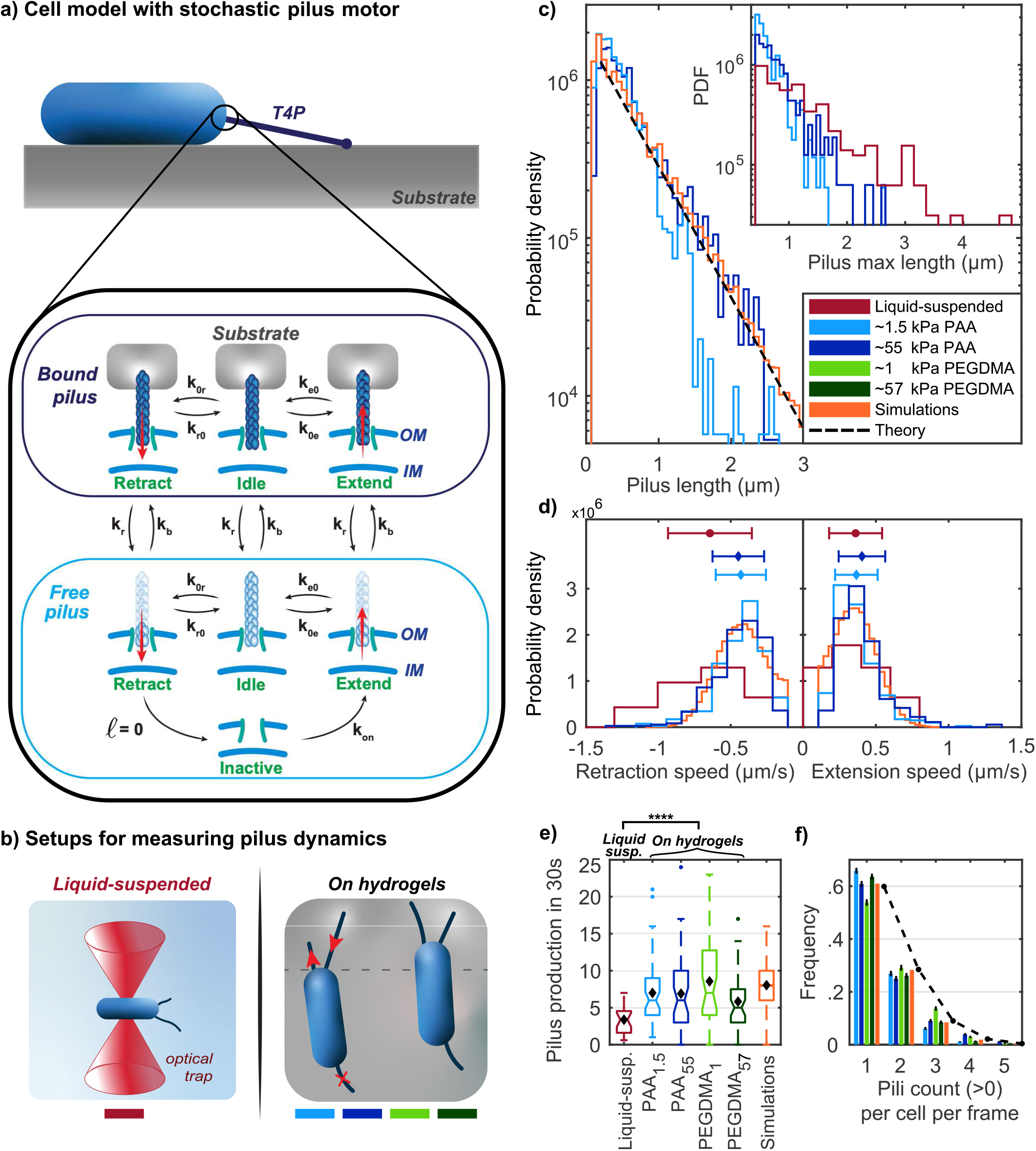
Systematic construction of a full model for T4P of bacteria on surfaces. a) Cell model and state diagram of the stochastic pilus-motor model. Different states describe pili that are bound to the substrate (dark blue) and pili that can extend and retract freely (light blue). b) Using an optical trap, cells are suspended in liquid for measuring only dynamics of free T4P as in Ref. [16]. Alternatively, T4P dynamics are quantified for cells lying on various substrates. c) T4P length distributions for different substrates. Theory predicts exponential distributions ∝ *e*^−*Kℓ*^, see main text. Inset: distributions of maximum length during full extension-retraction cycles. d) Distribution of average retraction and extension speeds per pilus. Error bars show the mean and the standard deviation. Simulated speeds are sampled and averaged over the time intervals that correspond to the frame rate in experiments. e) Number of T4P produced in 30 seconds. Black diamonds mark the mean of the data. Cells suspended in liquid produce significantly fewer T4P than cells on surfaces (*p* = 3.5 × 10^−9^, Welch’s t-test). f) Number of T4P per cell counted in individual images. Only cells with pili are considered. The histogrammed probability density is found by bootstrapping and error bars show the standard error of the mean. Orange line shows the theoretically predicted Poisson distribution. Simulation parameters model T4P on stiff PAA gels (*G*′ ≃ 55 kPa); see Supplementary text. See Supplementary Section 2.2 and Tab. S3 for data statistics.

Bacterial colonization depends on the physiochemical properties of the surface, see e.g., Refs. [23–25]. We therefore ask whether pilus activity depends on extracellular mechanical properties and quantify the T4P statistics on polyacrylamide (PAA) and polyethylene glycol-derived (PEGDMA) hydrogels of different stiffness. Comparison of results for soft and stiff hydrogels, with shear moduli of about 1 kPa and about 55 kPa respectively, however shows that substrate stiffness is not a systematic, major determinant of T4P statistics. While T4P on soft PAA gels are slightly shorter than on rigid PAA gels, a similar dependence on substrate stiffness is not found for PEGDMA gels, see Supplementary Fig. S4. Figure 2d shows the effect of surface interaction on pilus extension and retraction speeds. For cells on hydrogels, the speed distributions do not significantly depend on the type of substrate. However, T4P of cells on hydrogels have a significantly lower mean retraction speed than those of liquid-suspended cells. This finding is consistent with established force-velocity relationships for the T4P machinery, where mechanical loading of bound T4P reduces retraction speed [26].

Cell movement is not only determined by the dynamics of individual T4P, but also by their number. We find that, on average, bacteria on all substrates produce about twice as many pili per unit time as bacteria that are held in suspension, see Fig. 2e. This upregulation of T4P production is consistent with observations that transcription of the major pilin gene *pilA* is repressed for cells in suspension [27] and that polar FimW localization upon surface contact correlates with increased T4P production [28]. Figure 2f shows the numbers of simultaneously visible T4P per cell, counted for each image frame. For our conditions, *P. aeruginosa* rarely generates more than a single pilus simultaneously. Across hydrogels, no significant difference is found regarding the mean numbers of simultaneously measured T4P per cell. Pilus lifetime, defined as the time between appearance and complete retraction, is not significantly affected by substrate stiffness, see Supplementary Fig. S4g.

#### Model for free T4P

To explain the T4P statistics theoretically, we construct a model that is represented in the part of the state diagram labeled “free pilus” in Fig. 2a, see Methods Sec. I A. Thus, surface interaction is assumed to be rare, which is justified below. We also assume that T4P are retracted and extended with constant speeds *v*_*r*_ and *v*_*e*_, respectively. A fully retracted pilus can transition into a hidden, inactive state, from which it can re-appear stochastically. From these assumptions, we derive explicit formulae for T4P statistics, see Methods Sec. I A. The resulting distribution of the pilus length *ℓ* is

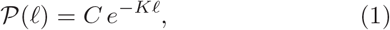

where the transition rate constants depicted in Fig. 2a result in a lengthscale 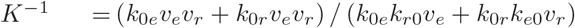 and *C* is a normalization constant. The measured pilus length distributions agree with the predicted exponential length distribution, see Fig. 2c. Moreover, the model predicts that the ratio of the probabilities to find an extending or retracting pilus only depends on the ratio of the processes’ speeds as *P*_e_*/P*_r_ = −*v*_*r*_*/v*_*e*_. Experimental data confirms this prediction with significance levels of [0.05 − 0.01], see Supplementary Tab. 1. For liquid-suspended cells, a fit of the analytical model confirms previously measured values of the rate parameters [16]. However, for cells on substrates, the average T4P are shorter than for liquid-suspended cells, see Fig. 2c, which cannot be explained in our model by the slower T4P retraction on surfaces. Based on a parameter variation, we conjecture that the shorter pili on substrates likely result from an increase of the parameter *k*_*e*0_, which determines the rate at which T4P switch from extension to idle. On a molecular level, this finding suggests that unbinding of the polymerization ATPase PilB is the major determinant of T4P length for bacteria on substrates.

The average lifetime of T4P, 𝒯_life_, determines a time scale during which pili can attach to their environment. We calculate 𝒯_life_ as mean first-passage time from emergence and to complete retraction, see Supplementary Section 2.2. The theoretical prediction is 𝒯_life_ = 2.64 s, which falls well within the range of lifetimes [2.2 − 4.0] s that we measure for different substrate conditions. Bootstrap analysis is employed for experimental data to take into account the occurrence of short, invisible pili during the finite image-recording intervals, see Supplementary Fig. S5. Our linear-rate-equation model implies that the numbers of T4P per cell must obey a Poisson distribution with a mean 𝒩_*p*_ = *k*_on_𝒯_life_. This prediction agrees well with the experimental results, see Fig. 2f. Measurement of the exponentially distributed waiting times between appearance of a new pilus yields *k*_on_ ∈ [0.25 − 0.4] s^−1^, see Supplementary Fig. S1.

#### Most T4P do not contribute to twitching

How do T4P coordinate extension-retraction dynamics and extracellular attachment to drive twitching migration? Recently, it has been suggested that individual *P. aeruginosa* T4P in an extended state are able to sense surface contact to initiate retraction [29]. To clarify whether such a mechanism plays a role for our cells, we perform a quantitative analysis of complete cycles of individual T4P-substrate interactions, including extension, substrate binding, retraction, and substrate unbinding. Since the unambiguous identification of the surface-binding events is challenging, we focus on the identification of those pili that visually form a mechanical connection with the surface and assume a tensed state during retraction, possibly displacing the bacteria. These pili are called “contributing pili”, see Fig. 3a.

**Figure 3.**
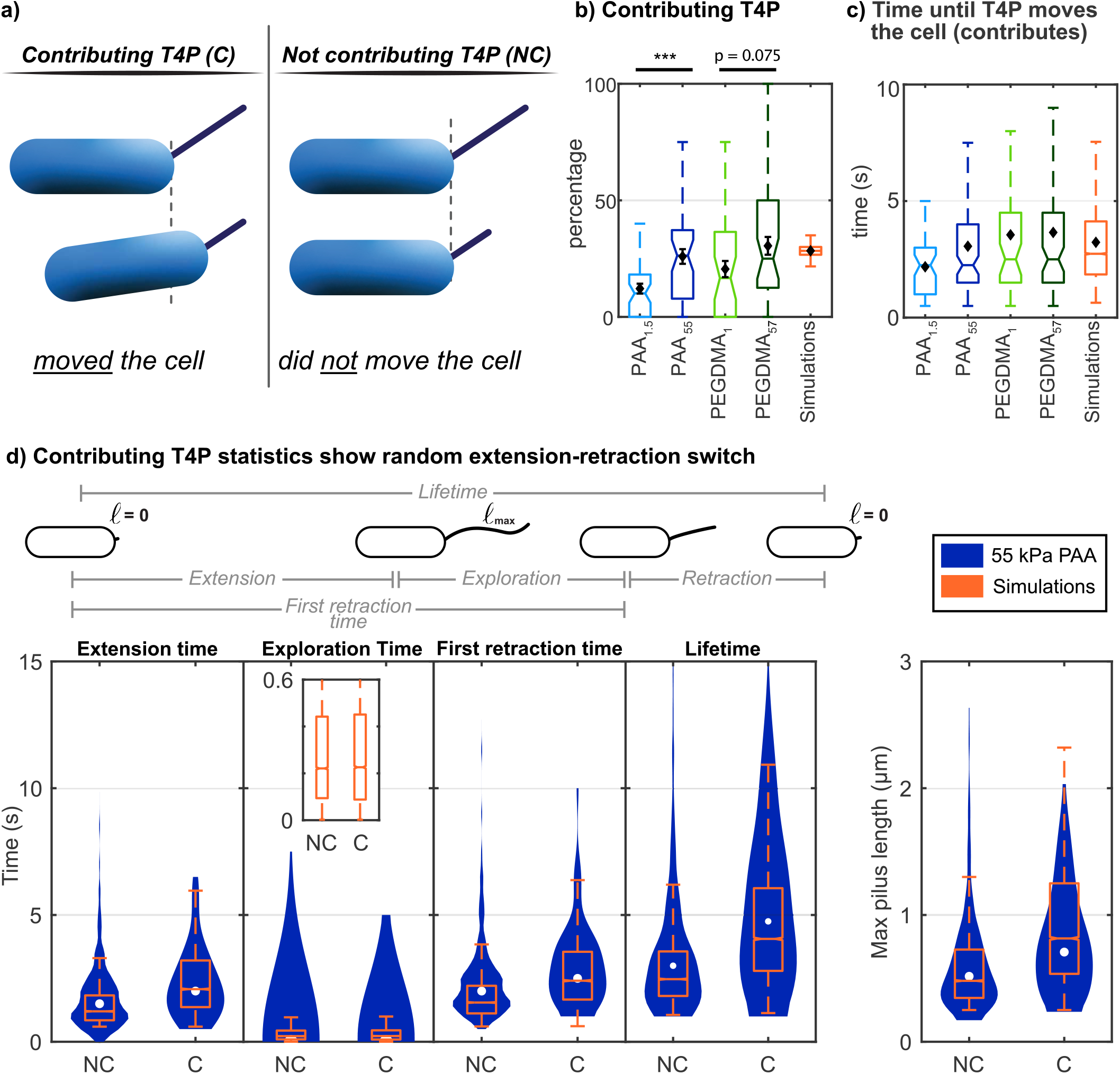
T4P life cycles and surface interaction can be explained without an attachment-dependent retraction trigger. a) Illustration of T4P categories. Only bound and tense T4P can contribute to twitching. b) Fraction of contributing T4P per cell are higher on stiffer substrates. Significance of having different medians is *p* = 0.00044 on PAA and *p* = 0.0753 on PEGDMA (U test). Error bars shows the standard error of the mean. c) Box plots of the time from creation of a pilus until the first time it is observed to be bound-and-tensed (contributing). d) Comparison of the statistics of contributing (C) and non-contributing (NC) pili. Blue violin plots show data from bacteria on 55 kPa PAA, where circles mark the median. Simulation results exclude pili that could not reach 0.25 *μ*m due to the attachment condition imposed. Inset: enlarged simulation results. See Supplementary Tab. S3 for data statistics.

We find that only 10% to 50% of pili actually contribute to twitching, see Fig. 3b. Both for PEGDMA and PAA hydrogels, the fraction of contributing pili increases with gel stiffness. This increase of contributing pili can be correlated with smaller hydrogel pore size and hence, a higher probability of substrate attachment. Pore sizes for PAA are in the range [10 − 40] nm [30] and [1 − 5] nm for PEGDMA gels [31]. Note that the pore size of commonly used 0.5 % agaraose gels is an order of magnitude larger, in the range of [500−1200] nm [32, 33]. Thus, T4P are expected to interact less frequently with agarose gels compared to more dense PAA or PEGDMA gels, which explains why T4P length statistics are similar for liquid-suspended cells and cells on 0.5 % agarose gels [16]. However, chemical substrate properties may affect the binding rate if the time scale set by diffusive encounter of a pilus binding site is not limiting the process.

To parametrize our model for T4P-substrate interaction, we next measure the time intervals between various events in contributing and non-contributing T4P. For example, substrate-binding rate constants are inferred from the average time between the emergence of a new pilus and its first contributing retraction, see Fig. 3c. We also test the hypothesis that an interaction between extended T4P and the substrate initiates a switch to the retraction state [29]. If this hypothesis were correct, contributing pili were expected to display 1) a shortened mean “exploration time”, i.e., a shortened mean time during which they wait in the fully extended state until retraction, and 2) a shortened time between pilus genesis and first retraction. Both predictions are not supported by the data. Figure 3d shows that the exploration times for contributing- and not-contributing pili are distributed similarly with exploration times ≤ 0.5 s in more than 80% of the events for all hydrogel conditions. Moreover, contributing pili on average have a larger first retraction time, i.e., start retracting later than not-contributing pili. This finding can be explained by the fact that contributing T4P are more likely to come from a T4P subpopulation that is long-lived and therefore more likely to bind to the substrate. For the same reason, contributing pili also spend a longer time on average for extension. In addition, the mean lifetime and length of contributing T4P is greater than that of non-contributing T4P, see Supplementary Fig. S9. Simulations show that this finding results from from a slower retraction of contributing pili due to their mechanical loading.

#### Passive adhesions reduce T4P-driven motion

Due to the low number of pili per cell, we rarely observe counteracting pili at both ends of bacteria that lead to a tug-of-war, as reported for the bacterium *Neisseria gonorrhoeae* [34, 35]. Rather, our time-lapse images suggest that the propulsion of bacteria through T4P retraction is obstructed by apparent cell-substrate forces without any detectable pili being present to hold the cells back. Furthermore, we observe cells that are only slightly displaced by the retraction of T4P at the leading pole and suddenly return to their original position after the pulling T4P detaches from the substrate, see Fig. 4a,b. These events, which we term “snap-back”, occur on elastic hydrogels. The presence of T4P-independent cell-substrate adhesions is also supported by computer simulations that show that cells with T4P at opposing poles do not produce the observed variation in migration dynamics, see Supplementary Fig. S8.

**Figure 4.**
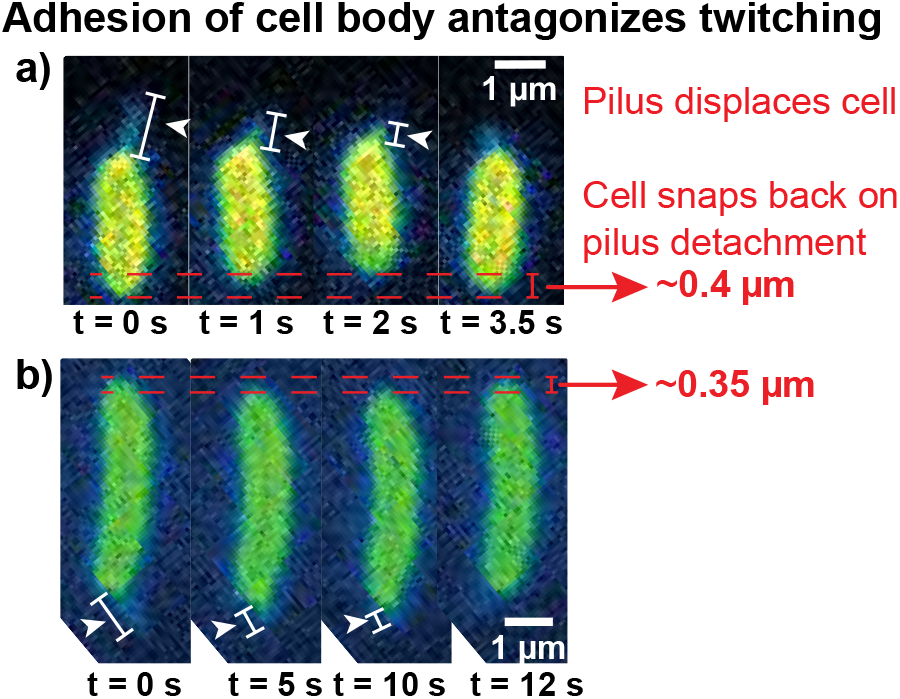
Snap-back events during twitching support a role of passive adhesions linking cell body and substrate. a) T4P pulls the cell forward without visible contribution of further T4P at the rear of the cell. Following disengagement of the T4P from the substrate, the cell instantly snaps back to its original position. b) Further example of a snap-back event.

### B. Twitching migration

#### Two regimes of random motion

Bacterial random walks are characterized by the ensemble-averaged mean-squared displacement (MSD) at time *τ*, denoted by Θ(*τ*), see methods. For migration of WT cells on short times, 1 s ≲ *τ* ≲10 s, we find that the mean-squared displacement scales as Θ(*τ*) ∝ *τ* ^*α*^ with *α* ≃ 1, similar to regular diffusion, see Fig. 5a. For long times, with *τ* in the range of minutes, we observe superdiffusive motion with *α* > 1, in the range [1.1 − 1.4], see Figs. 5a,b. Similar exponents for the long-time mean-squared displacement have been previously reported for *P. aeruginosa* twitching on glass [36, 37]. Of course, on very long time scales, diffusive behavior with *α* = 1 is expected, see Supplementary Fig. S3. The mean-squared displacements measured for migration of WT cells on four different hydrogels show qualitatively the same two regimes, although the magnitudes of the mean-squared displacements differ substantially. The observed mean-squared displacement can be faithfully reproduced in simulations where passive adhesions are modelled as stochastic bonds with a linearly elastic response.

**Figure 5.**
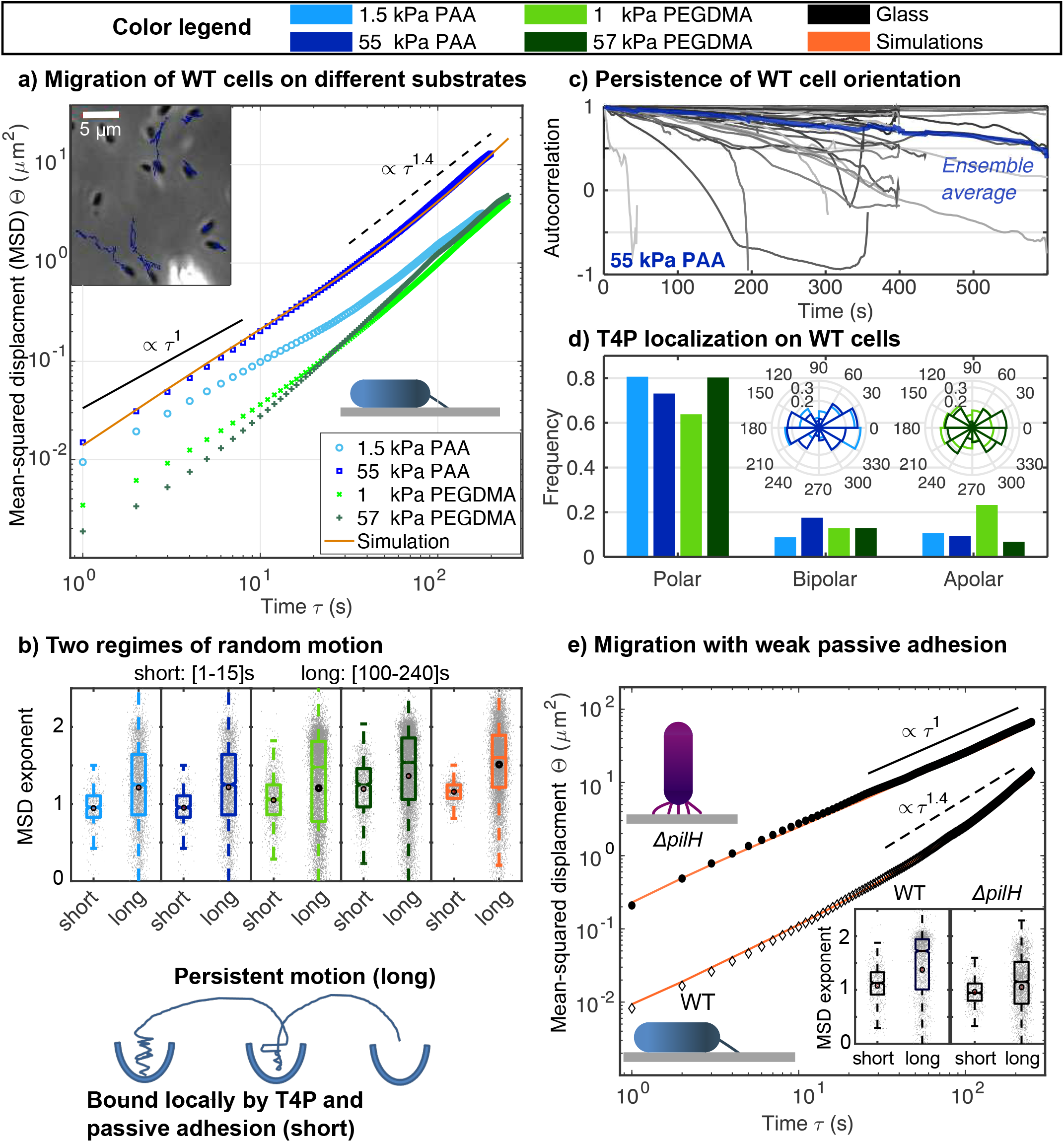
Pili and passive adhesions together determine two time scales of migration. a) Ensemble-averaged mean-squared displacements (MSDs) of wild-type (WT) cells on different substrates. b) Exponents of MSDs are different on short and long time scales. Dots show individual measurements. An exponent close to unity results from random motion of a cell that is locally bound before it explores other areas with persistent motion on longer time scales. c) Directional autocorrelation of cellular long axis decays over several minutes, see Supplementary Fig. S7 for the data regarding other gels. d) Statistics of T4P localization. At any time-point, a cell is labelled as polar if it has pili only on one pole, bipolar if there are pili on both poles, and “apolar” if it has pili protruding from the the sides of the cell and possibly also from the poles. d-inset) Angular distribution of pili with respect to the long axis of cells, which is chosen with random orientation, irrespective of the direction of motion. e) Ensemble-averaged MSDs of WT cells and Δ*pilH* mutants twitching on glass. Due to their predominantly vertical orientation on the substrate, Δ*pilH* mutants undergo random motion with an exponent that is only slightly larger than unity. Symbols represent the mean and error bars are standard deviations of the data.

#### Persistent motion on longer time scale

The superdiffusive character of the random walks can be explained by a slow change of the orientation of the bacterial long axis on the measurement time scale. Figure 5c illustrates that the autocorrelation of the angle of the long axis decays with a time scale around 15 minutes, see also Supplementary Fig. S7. Polar cells, with T4P only at one end, occur in [70 − 80] % of the data within sliding observation windows of 30 s, see Fig. 5d and Supplementary Fig. S2. Note that a change of the dominant pole, either spontaneously or through asymmetric division, occurs on time scales larger than 10 minutes and are therefore not frequently observable in our data. Cell polarity is approximately the same for all substrates and apolar emergence of pili from the long side of the cell body is rare. Simulations show that the experimentally quantified percentage of T4P on one pole is sufficient to explain superdiffusive motion on a time scale of 100 s ≲ *τ* ≲ 1000 s. Random motion with a persistent character can enable bacteria to follow potentially detected directional cues [27, 38].

#### Passive adhesions increase trapping on short time scales and persistence on long time scales

The regime of random motion with Θ(*τ*) ∝ *τ* found in the experiment for short times is also seen in simulations, where it occurs if passive adhesions transiently hold the cells at one spot. These are the snap-back events that are caused by the intermittency and varying spatial orientation of pilus-based force-generation, Fig. 4. Increasing the passive adhesion strength in the simulations leads to a broadening of the *α* ≃ 1 regime at short times. Simulations also show that the ratio of contributing pili correlates with the onset of persistent motion on longer time scales, where the higher the ratio of contributing pili, the earlier the onset of persistent motion.

Passive adhesions can also contribute to the persistence of motion by keeping the cell body pointing along one direction. To test experimentally how reduced persistence affects migration, we quantify the mean-squared displacement for hyperpilated Δ*pilH* mutants, see Fig. 5e, which predominantly adhere to a surface with an upright cell body, such that the long axis is oriented approximately normal to the surface. This orientation is likely caused by simultaneous activity of many pili on one pole, which produce a torque that lifts the body upwards [36, 39]. Upright standing presumably results in low passive adhesive forces since the fraction of the cell body in contact with the surface is reduced. As a result of the reduced persistence, the mean-squared displacement characterizing the random walk of Δ*pilH* mutants is well-described by a MSD exponent *α* that is only slightly larger than unity. Our simulations also show that the random walk of the Δ*pilH* mutants is significantly affected by fluctuations that do not result from pilus activity, which is not the case for wild-type cells that adhere passively to the substrate. Together, these findings suggest that passive surface-adhesion of the *P. aeruginosa* wild-type is essential for a migration dynamics that is reminiscent of random motion in a rugged potential landscape, where local valleys are explored on short time scales and large excursions occur on long time scales.

### C. Surface adhesion under shear flow

#### Twitching reduces surface-adhesion under shear

T4P are generally considered to be determinants of cell attachment. However, T4P retraction can also detach cells from the surface and it is therefore unclear whether twitching reduces or increases the residence time of bacteria under shear flow. To answer this question, we perform shear-flow experiments using a microfluidic setup and record cell detachments for different flow rates, see Fig. 6a. Figure 6c shows data for pilated wild-type cells that are classified as twitching and non-twitching. Both groups have T4P visible in the fluorescence channel and the average number of T4P per cell is slightly higher for twitching cells (*p* = 0.048 two-sample t-test), 1.27 ± 0.077 for twitching and 0.99 ± 0.117 for non-twitching cells. For all applied flow rates, we find that the ratio of detached cells after 20 minutes is is higher for the twitching bacteria. Thus, cell motion caused by T4P-activity correlates with reduced surface adhesion in our experiments.

**Figure 6.**
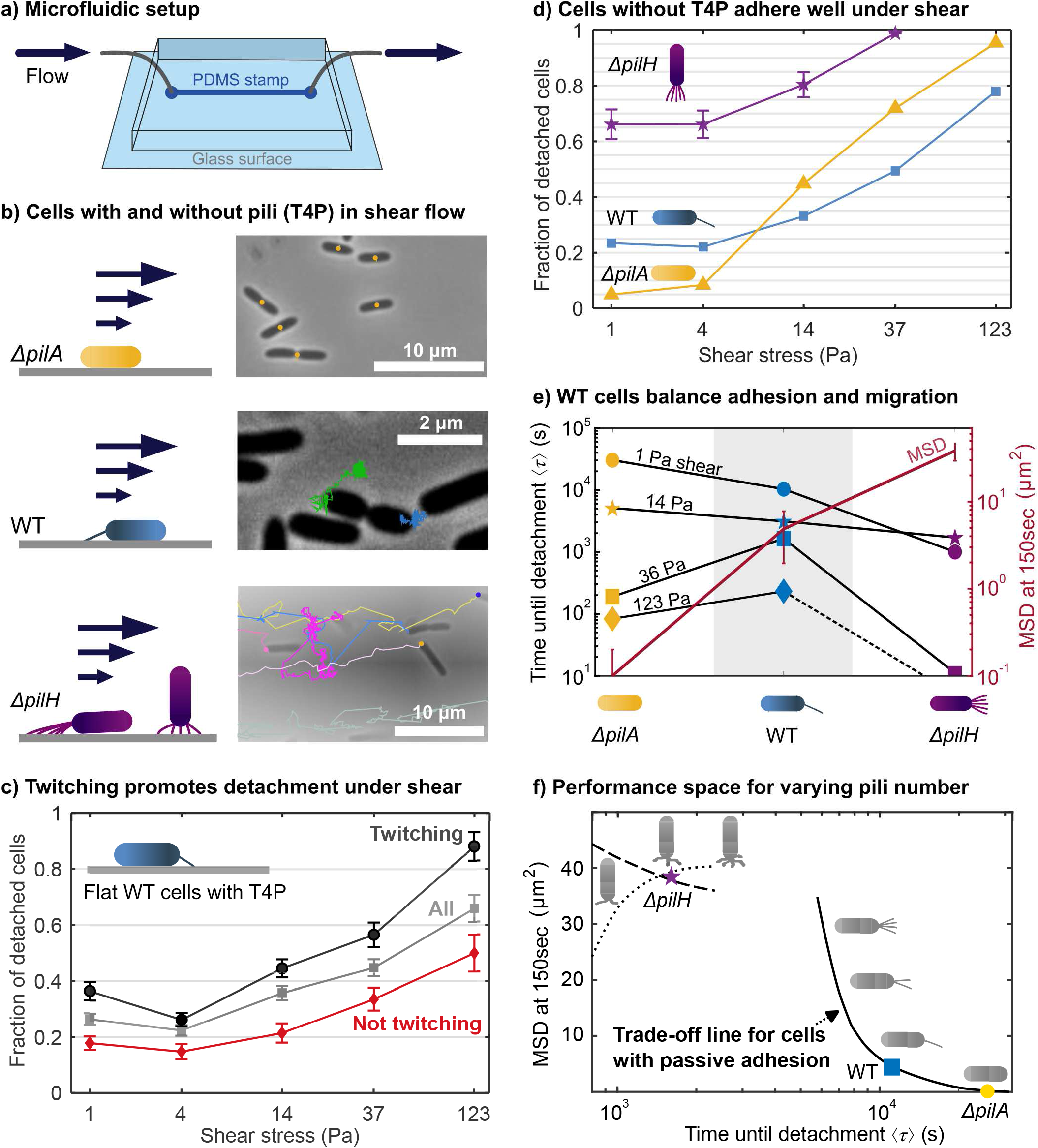
Balance of surface adhesion and migration. a) Setup of shear-flow experiments. b) Sample trajectories of three strains in shear flow with stresses of 1 Pa. b) Mean fraction of flat, adhering WT cells with T4P that detach under flow in 20 minutes. Fluorescence imaging of T4P is used to categorize cells as twitching or non-twitching prior to the onset of flow. Error bars show the standard error of the mean. c) Mean fraction of cells that detach under flow in 20 minutes. For Δ*pilH* mutants, detachment is quantified by counting attached cells every minute and error bars show the standard deviation of the detachment ratios for every minute mark. For WT cells and Δ*pilA* mutants, the standard errors of the mean detachment ratios are smaller than symbols in the plot. e) Time until surface detachment and MSD show opposite trends with increasing number of pili from Δ*pilA* to Δ*pilH* mutants. WT cells neither adhere best nor migrate best in weak shear, but achieve a trade-off that allows them to optimally combine both properties over a wide range of shear rates. MSD values are recorded in absence of flow and error bars show 95% confidence intervals. Error bars for ⟨*τ* ⟩ are smaller than the symbols. f) Qualitative structure of the adhesion-migration performance-space as a function of the pilus-generation rate. Simulation of detachment is done for 4 Pa shear stress and the MSD is recorded without flow. Line: flat cells with passive adhesion, dashed line: upright cells with noise enhancing cell diffusion, dotted line: upright cells at low noise. Passive adhesions enforce a trade-off. For upright walking cells, a simultaneous increase of MSD and detachment time may be possible for low noise and weak attachment.

#### Passive adhesions are essential for surface-adhesion under shear

To further clarify the role of T4P for surface adhesion under flow, we compare results for wild-type cells with mutants without T4P (Δ*pilA*) and hyperpilated cells (Δ*pilH*), see Fig. 6b,d. Typical shear stresses encountered by a bacterial pathogen in the human host host range from the order of 0.01 Pa in the proximal renal tubule [40] to the order of 10 Pa in vascular beds [41]. At these low shear stresses, the smallest fraction of detached cells is achieved by the Δ*pilA* mutant, which can only adhere passively. In contrast, Δ*pilH* cells which have many T4P are most easily detached. This result suggest that passive adhesions that anchor cells to a surface are by themselves more effective than T4P in preventing detachment under low shear stress.

For high shear stress above around 14 Pa, the lowest fraction of detached cells is achieved by the wild-type. Thus, having a few pili in addition to passive adhesions is the best way to stick under high shear conditions, see Fig. 6c. We suggest that the advantage of pilated cells under high shear results from a tightened horizontal alignment of the cell body with the surface, which promotes efficient engagement of passive adhesions, see Ref. [42]. Since pilated wild-type cells attach to the surface via one pole, hydrodynamic torque tends to align the cell body with the surface while torque produced by pilus retraction pulls the cells upwards. For a shear stress of 14 Pa, we estimate the hydrodynamic force that tilts the cells to be about 10 pN, which is comparable to the force magnitude generated by T4P [14]. Therefore, such shear stresses can be sufficient to maintain the cell body close to the substrate.

#### The adhesion-migration trade-off

To understand how *P. aeruginosa* combines surface attachment with migratory exploration, we compare the mean time until detachment under shear flow with the cells’ mean-squared displacement absence of flow, see Fig. 6e, Supplementary Figs. S11-S13. The two quantities show an opposite dependence on T4P numbers, where the time until detachment decreases as Δ*pilH* → WT → Δ*pilA* and the mean-squared displacement increases as Δ*pilA* → WT → Δ*pilH*. The crossing of the curves illustrates that wild-type cells combine the advantage of mutants having no T4P (good surface attachment, poor migration) and of mutants having many T4P (poor surface attachment, good migration).

Optimization can be visualized in a trait-performance space by plotting simulation results for detachment times against mean-squared displacements, see Fig. 6f. Since cells are found to generate more T4P on surface contact, see Fig. 2f, the T4P generation rate is assumed to be a parameter that cells can control for example by the Pil-Chp system [43, 44]. Accordingly, we vary the rate of T4P generation rate in our model. Detachment times are predicted qualitatively without taking hydrodynamics into account. For cells that generate passive adhesions, simulations reveal an inverse relationship between adhesion and migration. Thus, both quantities cannot be optimized simultaneously by wild-type cells and the curve can be interpreted as a Pareto front quantifying the trade-off between two incompatible traits. However, this front is not a continuous line since a high number of T4P leads to an upright cell configuration in experiments. Simulations of cells in upright configuration show that loose attachment and the resulting “Brownian-like” motion can allow both, an increase and a decrease of the mean-squared displacement with T4P numbers. Thus, the vertical configuration does not necessarily enforce a trade off between adhesion and migration. Nevertheless, the experimental results for the wild type are located on the right side of the discontinuity, perhaps even in the corner of the trade-off curve where the variances are large.

## II. DISCUSSION

Type-IV pili are a ubiquitous bacterial motility machinery and also an important virulence factor determining the severity of infections involving Pseudomonads [45]. We systematically dissect the stochastic dynamics of T4P when cells interact with surfaces and construct detailed mathematical models connecting the motor dynamics to migration of *P. aeruginosa*. Motor states, spatial pili organization, and binding events, are carefully characterized on the seconds-to-minutes time scale. We find that extension and retraction occur randomly. However, the average number of T4P increases on surfaces contact. Unlike the cytoskeleton of mammalian cells [46], we find no indication that regulation of the bacterial T4P machinery depends on substrate stiffness. For *P. aeruginosa*, we do not observe a tug-of-war between opposing T4P as seen for coccoid, peritrichously piliated *Neisseriae* [35], where sliding (kinetic) substrate friction reportedly affects motion, at least on the level of microcolonies [47]. Instead, our results are consistent with a model where Pseudomonads exhibit passive, sticking substrate friction that may result from physiochemical adsorption, fimbriae, and secreted adhesins such as Psl, Pel, and alginate exopolysaccharides [48–50]. Passive adhesions resist T4P-driven motion, trap cells locally on short time scales, and contribute to persistent motion by preventing random reorientation.

In shear flow, *P. aeruginosa* achieves a trade-off between fast migration and shear-resisting surface-attachment by maintaining a low average number of T4P. The number of T4P depends on a negative-feedback regulation of *pilA* transcription, as well as on the Chp chemosensory system, which includes the histidine kinase ChpA and two response regulators, PilG and PilH [27, 28, 51, 52]. Our results show that the production of T4P is upregulated on surface contact, but apparently only to a level where T4P activity does not abrogate passive surface adhesion. Thus, cells gain adhesiveness by sacrificing migration speed. Trade-offs between conflicting vital requirements are common in biological systems [53]. Examples include accuracy-efficiency [54] and growth-adaptability [55] problems, which can be solved on the level of individual cells or populations. The balance of adhesion and migration provides a new perspective to analyze motion optimality of diverse biological systems across scales, ranging from bacteria and algae in complex environments to immune cells and insects.

## III. METHODS

### A. Strains, growth conditions, and pilus labeling

Cells are grown and prepared for imaging as described previously [16]. In brief, the PilA cysteine knock-in mutant *P. aeruginosa pilA*-A86C is grown in liquid lysogeny broth (LB) Miller (Difco) in a floor shaker at 37^°^C. For imaging, Alexa488 maleimide dye (Fisher A10254) is suspended at a concentration of 2.5 mg/ml in anhydrous DMSO and stored −20^°^C. A fresh aliquot is used for each experiment to avoid freeze-thaw induced degradation of pilus labeling efficiency. Overnight grown cells are diluted 1:1000 in fresh LB medium and grown to mid log phase (OD600 = 0.4). Dye is added 1:100, incubated for 45 minutes at 37^°^C without shaking in the dark. Cells are washed twice in low autofluorescence EZ rich medium (Teknova) using a tabletop centrifuge at 8 krpm for 30 seconds and resuspended in 20 *μ*l fresh EZ rich medium.

### B. Hydrogel preparation

PAA gels [56] and PEGDMA gels [31] are made as described previously and mounted in a perfusion chamber (Warner Instruments). Prior to experiments, round, 40 mm coverslips (Fisherbrand) are methacrylate silanized by submerging plasma cleaned coverslips in 2% 3-(trimethoxysilyl) propylmethacrylate in 95% ethanol for 10 minutes. Hydrophobic round 12 mm coverslip (Fisherbrand) are prepared by submerging coverslips in Sigmacote (Sigma) for 10 minutes. All coverslips are subsequently rinsed three times in 100% ethanol and dried upright on Kimwipes. For preparation of gels, either 2% bis-acrylamide (BAA) and 40% polyacrylamide (AA) solutions (Bio-Rad) are diluted to 7.5% AA + 0.05% BAA (*G*′ ≃ 1.5 kPa, [57]) and 12% AA + 0.6% BAA (*G*′ ≃ 55 kPa, [57]) or PEGDMA (Mn 750, Sigma 437468) and 1.2M (356 mg/ml) 2-methacryloyloxyethyl phosphorylcholine (MPC, Sigma 730114) are diluted to 1.1% PEGDMA + 50% MPC (*G*′ ≃ 1 kPa, [31]) and 6.3% PEGDMA + 50% MPC (*G*′ ≃ 57 kPa, [31]) in 496.5 *μ*l ddH2O, respectively. These solutions are degassed for 10 minutes in a vacuum chamber. To initiate polymerization, 2.5 *μ*l of 10% Ammonium persulfate (APS, Bio-rad) and 1 *μ*l Tetramethylethylenediamine (Temed, Bio-rad) are added and gently mixed by pipetting. Hydrogels are then allowed to polymerize between two chemically modified coverslips that were prepared prior to experiments. 10 *μ*l of unpolymerized gel is added between both coverslips resulting in an approximately 50 *μ*m tall gel. Gels are polymerized for 30 minutes at room temperature, followed by removal of the hydrophobic coverslip. Finally, gels are washed three times and then incubated for 30 minutes in EZ rich medium.

### C. T4P dynamics on hydrogels

#### a. Imaging

To prepare experiments, excess liquid medium is carefully removed from the gel and the coverslip and 20 *μ*l of cell suspension is added. The inlets to the perfusion chamber are filled with EZ rich medium and the chamber is subsequently assembled carefully to avoid air bubbles in the system. The chamber is then mounted on a Nikon TiE microscope housed in an environmental chamber at 37^°^C. To remove unbound cells, a slight flow using a flow pump is introduced. Fluorescent pilus images are taken every 0.5 seconds using a white light LED source (Excelitas), a GFP filter set, a Nikon Plan Apo 60X WI DIC N2 lens, and an Andor Clara CCD camera with x1.5 zoom. Brightfield images of bacteria on gel are also taken with a 40X objective and the same setup.

#### b. Pilus data acquisition

Prior to analyzing the fluorescence images, duplicate images are employed to construct lookup tables for contrast optimization. Pili are then identified visually by comparing original and the processed images (the brightness and contrast of the images are adjusted automatically and despeckled by ImageJ, and displayed with a lookup table.). The tip and the base on the cell membrane are marked and recorded for each pilus, where the expected maximum measuring error is 1 pixel (0.07 *μ*m). The process is repeated for all bacteria with observable pili. The pili that are idle for the duration of the experiments are excluded from the analysis.

#### c. Cell tracking

The bacteria are tracked from phase-contrast movies semi-automatically with the ImageJ plugin MicrobeJ [58]. The plugin supports sub-pixel registration and the magnitude of tracking errors is distributed exponentially with a mean of 0.25 pixel lengths. First, the plugin segments and tracks automatically. Afterward, the segmentation and tracking errors are corrected by hand. Cells that are only partially adhered were excluded from the analysis.

### D. Flow experiments

#### a. Microfluidic flow chamber preparation

Microfluidic devices are made as described previously [59]. Briefly, soft lithography masks are designed in AutoCAD and printed by CAD/Art Services. A mold is then made on a silicon wafer (University Wafer) by spin coating SU-8 photoresist (MicroChem). Microfluidic chips are made by pouring freshly mixed and vacuum degassed polydimethylsiloxane (PDMS, Sylgard 184, Dow) over the mold. PDMS is cured for 2 hours at 60^°^C. Individual chips are cut out and bonded to plasma-cleaned 22 × 40 mm coverslips (Fisherbrand) for 1 hour at 60^°^C. Each chip has 6 parallel channels with width *W* ≃ 600 *μ*m and height *H* ≃ 110 *μ*m high or 55 *μ*m, and 2 cm length. Only one channel is used per experiment. The wall shear stress is estimated from the known flow rate as in Ref. [60].

#### b. Imaging and analysis

Flow experiments are performed on the same microscope as described above using a 100x oil immersion objective and 1.0x zoom. Flow chambers are filled with labeled cells and incubated for 15 minutes on the microscope at 37^°^C prior to starting the experiment. First, pili are imaged every second for 30 seconds with a maximum flow of 3 *μ*l*/*min. Second, cells are imaged every 1 second for 20 minutes in brightfield at flow rates of [25, 75, 250, 750, 2500] *μ*l*/*min for 55 *μ*m high channels with WT cells and and [100, 300, 1000, 3000, 10000] *μ*l*/*min for 110 *μ*m high channels with mutants. Due to the volume limit of the syringe pump, cells in the largest flow at stesses of 123 Pa could only be recorded for about 10 minutes. Cell detachment is quantified by counting the number of cells in each frame of a movie. Then, stretched exponential functions are fitted to the remaining fraction of cells [60]. Note that the resulting detachment statistics ignore diverse cell behaviors inside the field of view, such as sliding of the Δ*pilA* mutants along with the flow and random jumps of the Δ*pilH* mutants.

### E. Analytical model for free pili

The free-pilus model depicted in the inset of Fig. 2a yields the probability distributions *P*_…_(*ℓ, t*) for finding a pilus with length *ℓ* at time *t* in different states. The equations determining the probability distributions are are

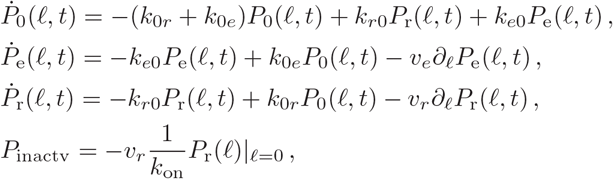

where the time-derivatives 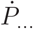 are assumed to vanish identically since we only require the steady-state solutions *P*_…_(*ℓ*). Using Kramers’ method for mean first-passJage times, the average T4P lifetime results as 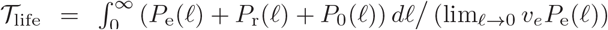. The number of T4P per cell, 𝒩_*p*_, obeys a production-degradation subprocess as 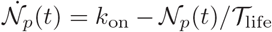. See Supplementary Section 2 for details.

### F. Simulation model for twitching cells

Cells are modelled as point-like particles having several T4P and passive adhesions. The equation of motion for the position x in a Cartesian, two-dimensional system at time *t* reads

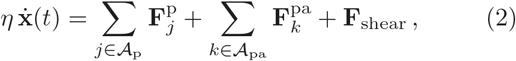

where 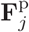 is the tensional force exertered by the *j*^th^ attached T4P (*j* ∈ A_p_), 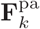 is the force transmitted by the *k*^th^ attached passive adhesion (*k* ∈ 𝒜_pa_), **F**_shear_ is the hydrodynamic shear force, and *η* is a friction constant. To avoid a modelling of unknown, complex hydrodynamics during cell detachment, the time until detachment under shear was taken to be proportional to the mean time until all cell-substrate bonds are broken in the simulations.

The unit vector **e**_*j*_ denotes the direction of the *j*^th^ pilus. The force transmitted by this pilus is given by 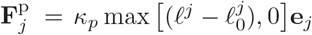 where *κ*_*p*_ is a spring constant, *ℓ*^*j*^ is the actual pilus length, and 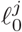 is its rest length. Due to visible filament buckling, T4P are assumed to transmit only tensional forces. Measurable T4P can be extending (e), idle (i), or retracting (r). In these states, the rest length changes with speeds *v*_e_ *>* 0, *v*_i_ = 0, and 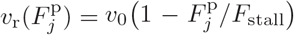, where *v*_0_ *<* 0. All pili with length 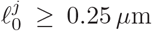 can independently bind to the substrate with rate constant *π*_*p*_. Pilus detachment occurs with a force-dependent rate 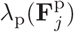. New pili are generated with rate constant *k*_on_. On first generation of a T4P, **e**_*j*_ is drawn from the measured orientation distribution, see Fig 5d. The direction vectors of attached pili are updated at every simulation time step based on the motion of the bacterium. When a retracting pilus reaches 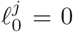, the pilus is deactivated and detached from the substrate.

Passive adhesins act as elastic anchors holding the bacterium to the surface, to which they bind with rate constant *π*_*pa*_. An attached passive adhesion bond transmits the force 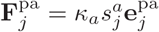 where 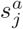 is its stretch. The orientation of the passive adhesin 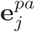 is initially taken to be antiparallel to the cell’s migration direction and is updated as the cell moves ahead. Breakage of passive adhesion bonds occurs with force-dependent rates 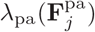. The number of passive adhesins that can bind the to the surface is assumed to be constant. For simulation of Δ*pilH* mutants, the number of passive adhesins is set to zero and an approximately isotropic pilus orientation is assumed. For Δ*pilH* mutants, white Gaussian noise is added to the trajectory of cells that are not attached.

## DATA AVAILABILITY

Simulation data that is generated and analysed during the current study is available from the corresponding author upon reasonable request.

## CODE AVAILABILITY

Code for the simulation of bacteria is available from the corresponding author on reasonable request.

## AUTHOR CONTRIBUTIONS

M.D.K. and B.S. carried out the experiments. A.N.S. and B.S. conducted the theoretical analysis. All authors contributed to the interpretation of the data and to the manuscript preparation.

## ACKNOWLEDGEMENTS

BS and ANS acknowledge funding by the European Research Council (g.a.No. 852585) and by the German Research Foundation (SPP 2332, SA 2643/3-1). ANS acknowledges support by the International Helmholtz Research School of Biophysics and Soft Matter (IHRS BioSoft). This work was supported by grant K22AI151263 from the National Institutes of Health to J.E.S.

