## Supplementary text and figures for "Type-IV pili tune an adhesion-migration trade-off during surface colonization of *Pseudomonas aeruginosa*"

May 7, 2023

### 1 Data Analysis

#### 1.1 Analysis of pilus dynamics

To calculate the retraction and extension speeds, the pilus lifetime is divided into two parts: before and after it reaches the maximum length. The average retraction and extension speed per pilus are calculated while excluding speed values smaller than 0.105 $\mu$m/sec (1.5 pixel/sec), thus excluding phases where pili are idle. T4P that are idle during most of the observation time are omitted from the analysis.

Table S1: Experimental data confirms the predicted relationship $-\langle v_r\rangle/\langle v_e\rangle \approx \langle P_e\rangle/\langle P_r\rangle$. The ratio of average extension-retraction velocities is compared to the ratio of average times spent in the extension-retraction states. Data includes only T4P with completely observed extension-retraction cycles.

| Gel | Ratio | All | C | NC |
|---|---|---|---|---|
| $PAA_{1.5}$ | $-\langle v_r\rangle/\langle v_e\rangle$ | 1.1585 | 1.1523 | 1.1591 |
| | $\langle P_e\rangle/\langle P_r\rangle$ | 1.1401 | 1.1012 | 1.1455 |
| $PAA_{55}$ | $-\langle v_r\rangle/\langle v_e\rangle$ | 1.0608 | 0.9386 | 1.1231 |
| | $\langle P_e\rangle/\langle P_r\rangle$ | 1.0414 | 0.9417 | 1.0737 |
| $PEGDMA_{1.5}$ | $-\langle v_r\rangle/\langle v_e\rangle$ | 1.1518 | 1.0299 | 1.1820 |
| | $\langle P_e\rangle/\langle P_r\rangle$ | 1.1611 | 1.1659 | 1.1603 |
| $PEGDMA_{55}$ | $-\langle v_r\rangle/\langle v_e\rangle$ | 1.1045 | 1.0142 | 1.1570 |
| | $\langle P_e\rangle/\langle P_r\rangle$ | 1.1048 | 1.0009 | 1.1387 |

#### 1.2 Pili distribution on the cell body

For each cell from which T4P data are collected, the positions of the poles are manually marked to determine the angle between the pili and the long axis of the cell. The

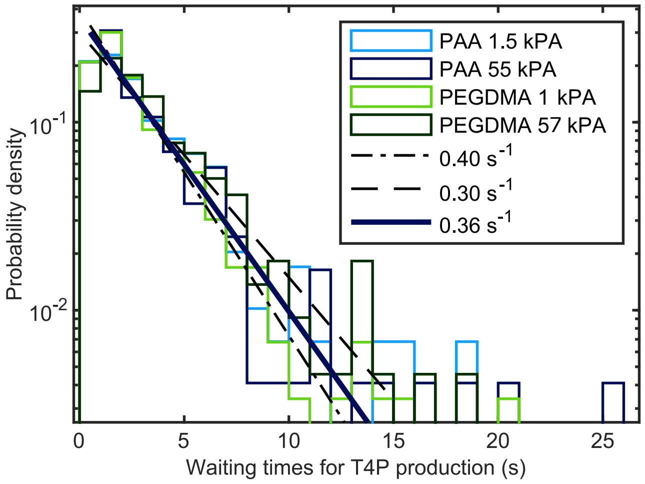


Figure S1: Distribution of time-interval lengths between the production of individual new T4P, measured cell by cell.

angle distributions shown in Fig. 5d of the main text are measured when the pili are at their maximum length. If a pilus originates within a radius of 10 pixel lengths around a cell pole, the pilus is said to originate from that pole. If the pilus does not meet this criterion for both poles, it is marked as apolar. Polarity is assessed for each time point in the image sequences. Cells are categorised as polar, bipolar, or apolar if pili emerge from one pole, both poles, or neither pole, respectively. Note that photobleaching limits the time period during which T4P can be observed, making it difficult to quantify the switching of the dominant cell pole by direct observation of pili.

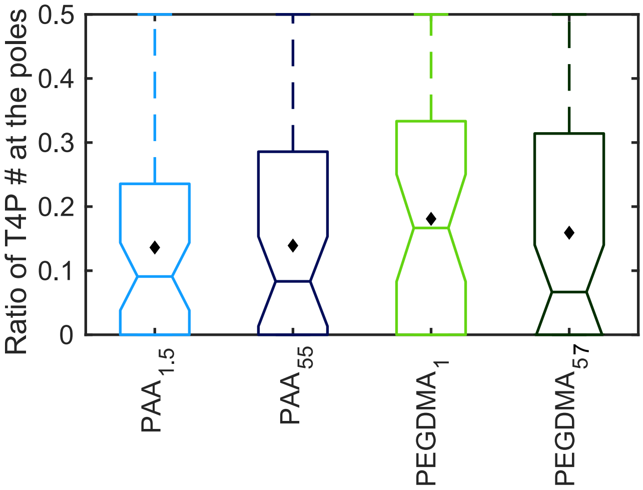


Figure S2: The number of T4P created at the trailing pole of a cell divided by the total number of T4P produced by a cell at both poles within 30 seconds (60 frames). The trailing pole is defined in this figure as the pole that produces fewer T4P. Zero means that there are no pili at the trailing pole. A value of 0.5 would imply equally many T4P at both poles.

##### 1.3 Cell-orientation autocorrelation

For each bacterium with index $i$, we denote the position of the two poles as $\mathbf{r}_1^i(t)$ and $\mathbf{r}_2^i(t)$, with the order of the poles being arbitrary for this analysis. A director is given by $\mathbf{e}^i(t) = (\mathbf{r}_2^i(t)-\mathbf{r}_1^i(t))/||\mathbf{r}_2^i(t)-\mathbf{r}_1^i(t)||$. Then, directional autocorrelation at lag time $\tau$ is given by

$$\mathcal{C}_{\text{dir}}^i(\tau) = \langle \mathbf{e}^i(\tau+t)\cdot\mathbf{e}^i(t)\rangle_t\,, \tag{1}$$

and the ensemble-averaged directional autocorrelation is $\mathcal{C}_{\text{dir}}(\tau) = \sum_i^{N_\tau}\mathcal{C}_{\text{dir}}^i(\tau)/N_\tau$.

##### 1.4 Mean-squared displacement

For a each bacterium with index $i$, we calculate a time-averaged mean-squared as

$$\Theta_i(\tau) = \left\langle \left(\mathbf{r}_x^i(\tau+t)-\mathbf{r}_x^i(t)\right)^2 + \left(\mathbf{r}_y^i(\tau+t)-\mathbf{r}_y^i(t)\right)^2\right\rangle_t \tag{2}$$

where $\tau$ is the lag time. The ensemble-averaged mean squared displacement for $N$ bacteria with varying trajectory length is calculated by

$$\Theta(\tau) = \sum_i^{N_\tau}\frac{\Theta_i(\tau)}{N_\tau}\,, \tag{3}$$

where $N_\tau \leq N$ is the number of bacteria that have been tracked for more than $2\tau$ time steps. To avoid fictitious jumps in the MSD that result from the different lengths of the trajectories combined with the cell-to-cell variability of migration, we calculate the increment of the ensemble-averaged MSD $\Delta\Theta(\tau) = (\Theta(\tau)-\Theta(\tau-1))/\tau$ and plot the integral of $\Delta\Theta(\tau)$ in the figures pertaining to the main text.

##### 1.5 Detachment under shear flow

For the data shown in Fig. 6c of the main text, piliated bacteria are classified as twitching or non-twitching based on the observation of pili-driven twitching in the fluorescence channel before the onset of flow and in the first few frames of the brightfield movies. The time at which bacteria detach from the surface is recorded. Bacteria that do not lie flat on the surface and bacteria without pili are excluded. Note that sometimes bacteria will detach but reattach a few microns away. Such an event is not counted as detachment.

The mean time until detachment shown in Fig. 6e-f of the main text is determined by counting the remaining cells in every time frame of an image sequence after onset of the flow. As suggested in Ref. [1], the fraction of cells remaining until time $t$ is fitted with a stretched exponential function. A brief time period $t_0$ is assumed to be required for the build-up of stationary flow after turning on the syringe pump ($t_0 \sim 100$s ). For $t \geq t_0$, we fit the remaining fraction of cells with $a\exp[-((t-t_0)/\tau)^\alpha]$, where $a$ and and $\alpha \in [0,1]$ are parameters and $\tau$ represents a timescale of survival see Figs. S11,S12, and S13. The mean time until detachment is calculated as $\langle\tau\rangle = \tau\Gamma(1/\alpha)/\alpha$ with $\Gamma(..)$ being the Gamma function.

##### 1.6 Representation of box plots

Bottom and top edges of the box indicate 25th and 75th percentiles, the central mark indicates the median with notches showing its 5% significance level (median $\pm$ $1.57(\text{q}_{75}-\text{q}_{25})/\sqrt{\text{n}_{\text{sample}}}$). Black diamonds always display the mean of the data and their error bars show the standard error of the mean. Dots represent the outliers that are more than 1.5 times the interquartile range away from the bottom or top of the box. Outliers above the limits of y-axis are not shown to clearly display significant statistics. Outliers of the simulations are not displayed for easier comparison of the statistically significant data.

#### 2 Analytical model for free T4P

##### 2.1 Model and steady-state solution

We propose a stochastic model to describe the retraction and elongation dynamics of pili, see the inset "Free pilus" in Fig. 2a of the main text. At any given time, a pilus site can either be active or inactive. An inactive site ($\mathbf{P}_{\text{inactv}}$) has no activity and no pilus. A pilus site stochastically switches to the active state with rate constant $k_{\text{on}}$. An active pilus site can either be elongating, retracting, or idling. Elongating T4P increase their length with speed $v_e > 0$. Retracting T4p reduce their length with speed $v_r < 0$). If a pilus is idling, the pilus length does not change until the motor state changes. The steady-state probability densities for pili with length $\ell$ in the elongating, retracting, and idling state are denoted by $\mathbf{P}_{\text{e}}(\ell)$, $\mathbf{P}_{\text{r}}(\ell)$ and $\mathbf{P}_0(\ell)$, respectively. A pilus site becomes inactive when the pilus is completely retracted. We assume that a pilus site has to go through the idle state to switch between the retraction and the elongation states. The steady-state probabilities are determined by

$$0 = -(k_{0r} + k_{0e})\mathbf{P}_0(\ell) + k_{r0}\mathbf{P}_{\mathrm{r}}(\ell) + k_{e0}\mathbf{P}_{\mathrm{e}}(\ell)\,, \quad (4)$$
$$0 = -k_{e0}\mathbf{P}_{\mathrm{e}}(\ell) + k_{0e}\mathbf{P}_0(\ell) - v_e\mathbf{P}'_{\mathrm{e}}(\ell)\,, \quad (5)$$
$$0 = -k_{r0}\mathbf{P}_{\mathrm{r}}(\ell) + k_{0r}\mathbf{P}_0(\ell) - v_r\mathbf{P}'_{\mathrm{r}}(\ell)\,, \quad (6)$$
$$\mathbf{P}_{\mathrm{inactv}} = -v_r\frac{1}{k_{\mathrm{on}}}\mathbf{P}_{\mathrm{r}}(\ell)|_{\ell=0}\,, \quad (7)$$

where $\mathbf{P}'_{\{\mathrm{e,r}\}}$ are derivatives with respect to pilus length $\ell$. Substitution of Eqs. (5),(6) into Eq. (4) yields the basic relation

$$v_e\mathbf{P}'_{\mathrm{e}}(\ell) = -v_r\mathbf{P}'_{\mathrm{r}}(\ell) \quad (8)$$

from which, by integrating out $\ell$, we obtain

$$v_e\mathbf{P}_{\mathrm{e}}(\ell)|_{\ell=0} = -v_r\mathbf{P}_{\mathrm{r}}(\ell)|_{\ell=0}. \quad (9)$$

Furthermore, using Eq (9) and Eq (4), we can employ for $\mathbf{P}_{\mathrm{e}}$, $\mathbf{P}_{\mathrm{r}}$, and $\mathbf{P}_0$ an Anatz $\propto e^{-K\ell}$ where the proportionality constant depends on $\mathbf{P}_{\mathrm{inactv}}$ and the other parameters. By substitution one finds the exponent in our Ansatz from Eq (5) as

$$K = \frac{k_{0e}k_{r0}v_e + k_{0r}k_{e0}v_r}{v_e v_r\,(k_{0e} + k_{0r})}\,. \quad (10)$$

Solutions for the steady-state probability densities in terms of the parameters and $\mathbf{P}_{\mathrm{inactv}}$ are given by

$$\mathbf{P}_0(\ell) = e^{-K\ell}\frac{k_{\mathrm{on}}\mathbf{P}_{\mathrm{inactv}}(k_{e0}v_r - k_{r0}v_e)}{v_e v_r(k_{0e} + k_{0r})}\,, \quad (11)$$
$$\mathbf{P}_{\mathrm{e}}(\ell) = e^{-K\ell}\frac{k_{\mathrm{on}}\mathbf{P}_{\mathrm{inactv}}}{v_e}\,, \quad (12)$$
$$\mathbf{P}_{\mathrm{r}}(\ell) = e^{-K\ell}\frac{k_{\mathrm{on}}\mathbf{P}_{\mathrm{inactv}}}{-v_r}\,. \quad (13)$$

Thus, independent of the non-obervable inactive state, the steady-state probabilities are connected by the following relations

$$\mathbf{P}_{\mathrm{e}} = -\frac{v_r}{v_e}\mathbf{P}_{\mathrm{r}}, \qquad \mathbf{P}_0 = \mathbf{P}_{\mathrm{r}}\frac{k_{r0}v_e - v_r k_{e0}}{v_e(k_{0r} + k_{0e})}\,. \quad (14)$$

Finally, the steady-state probability for the inactive state is calculated from normalization

$$\mathbf{P}_{\mathrm{inactv}} + \int_0^\infty (\mathbf{P}_{\mathrm{e}}(\ell) + \mathbf{P}_{\mathrm{r}}(\ell) + \mathbf{P}_0(\ell))\,d\ell = 1. \quad (15)$$

The full expressions for $\mathbf{P}_{\mathrm{inactv}}$ the other probability densities are given in Eqs. (16)-(19). The directly measurable probability density $\mathcal{P}(\ell)$ to find a pilus, with length $\ell$, in any observable state is provided in (20).

#### 2.2 Pilus lifetime

The analytical model naturally yields an average pilus lifetime via the auxiliary problem of mean-first-passage times. By means of Kramers' method we can calculate the mean first passage time, $T_{\mathrm{life}}$, for a pilus to start at $\ell = 0$ from the extension state (e) and end again in the completely retracted state, $\ell = 0$, in (r), which is the mean observable pilus lifetime. The flux boundary condition imposed on the extension state at vanishing T4P length is

$$J = \lim_{\ell\to 0} v_e\mathbf{P}_{\mathrm{e}}(\ell)\,. \quad (21)$$

Using the average occupancy of the states, Eq (20), Kramers' method allows one to calculate the mean pilus life time as

$$\mathcal{T}_{\mathrm{life}} = \int_0^\infty \mathcal{P}(\ell)d\ell \Big/ J = \frac{v_r(k_{0e} + k_{0r} + k_{e0}) - v_e(k_{0e} + k_{0r} + k_{r0})}{k_{0e}k_{r0}v_e + k_{0r}k_{e0}v_r}\,. \quad (22)$$

A direct measurement of the predicted exponential lifetime distribution is challening because many of the T4P are very short lived, with a lifetime less than a second. Such small and short-lived pili are hard to quantify given the spatial and temporal resolution limits of the microscopy data. However, simulation results for the parameter values listed in Tab. S2 match the measured data well when short lived T4P that occur for $< 1$ s in the simulations are omitted, see the orange box in Fig. S4g. The analytical formula for the average pilus lifetime yields $\mathcal{T}_{\mathrm{life}} \simeq 2.6$ sec for the chosen parameter values.

#### 2.3 Average number of T4P per cell

We assume a high number of inactive T4P transmembrane complextes [2] and that the states of individual pili are independent of each other. New pili are produced with an activation rate of $k_{\mathrm{on}}$. If the mean life time of pili is $\mathcal{T}_{\mathrm{life}}$, then the "average death rate" or full-retraction rate is $k_{\mathrm{death}} = 1/\mathcal{T}_{\mathrm{life}}$. The probability $\mathbf{p}(n,t)$ to find $n$ pili at time $t$ in any given cell obeys a master equation as

$$\dot{\mathbf{p}}(0,t) = k_{\mathrm{death}}\mathbf{p}(1,t) - k_{\mathrm{on}}\mathbf{p}(0,t)\,,$$
$$\dot{\mathbf{p}}(n,t) = k_{\mathrm{death}}(n+1)\mathbf{p}(n+1,t) + k_{\mathrm{on}}\mathbf{p}(n-1,t) - (nk_{\mathrm{death}} + k_{\mathrm{on}})\mathbf{p}(n,t) \ \text{ for n} > 0\,. \quad (23)$$

Using this resulting probability distribution, the average number of pili can be calculated as $\mathcal{N}_p(t) = \sum_{n=0}^\infty n\mathbf{p}(n,t)$. Algebraic manipulation of Eq. (23) yields an equation for the evolution of the average number of pili as

$$\dot{\mathcal{N}_p}(t) = k_{\mathrm{on}} - k_{\mathrm{death}}\mathcal{N}_p(t). \quad (24)$$

Supposing that the bacterium initially has no pili, one obtains

$$\mathcal{N}_p(t) = \frac{k_{\mathrm{on}}}{k_{\mathrm{death}}}\left(1 - e^{-k_{\mathrm{death}}t}\right)\,, \quad (25)$$

which, in the long-time limit converges to $\mathcal{N}_p \approx \frac{k_{\mathrm{on}}}{k_{\mathrm{death}}}$, i.e.,

$$\mathcal{N}_p \approx k_{\mathrm{on}}\frac{v_r(k_{0e} + k_{0r} + k_{e0}) - v_e(k_{0e} + k_{0r} + k_{r0})}{k_{0e}k_{r0}v_e + k_{0r}k_{e0}v_r}\,. \quad (26)$$

Also, the production-degradation process, Eq. (23), yields a steady-state solution $p^s(n)$ that obeys a Poisson distribution as

$$p^s(n) = \frac{1}{n!}\left(\frac{k_{\mathrm{on}}}{k_{\mathrm{death}}}\right)^n e^{-\frac{k_{\mathrm{on}}}{k_{\mathrm{death}}}}\,. \quad (27)$$

$$\mathbf{P}_{\text{inactv}} = \frac{k_{0e}k_{r0}v_e + k_{0r}k_{e0}v_r}{v_e k_{r0}(k_{0e} - k_{\text{on}}) + k_{e0}v_r(k_{0r} + k_{\text{on}}) - k_{0e}k_{\text{on}}(v_e - v_r) - k_{0r}k_{\text{on}}(v_e - v_r)}, \tag{16}$$

$$\mathbf{P}_0(\ell) = \frac{k_{\text{on}}(k_{r0}v_e - k_{e0}v_r)(k_{0e}k_{r0}v_e + k_{0r}k_{e0}v_r)\exp\left(-\ell\frac{k_{0e}k_{r0}v_e + k_{0r}k_{e0}v_r}{k_{0e}v_e v_r + k_{0r}v_e v_r}\right)}{v_e(k_{0e}(k_{\text{on}} - k_{r0}) + k_{\text{on}}(k_{0r} + k_{r0}))) - v_e v_r(k_{0e} + k_{0r})(v_r(k_{\text{on}}(k_{0e} + k_{0r} + k_{e0}) + k_{0r}k_{e0})}, \tag{17}$$

$$\mathbf{P}_{\text{e}}(\ell) = \frac{k_{\text{on}}(k_{0e}k_{r0}v_e + k_{0r}k_{e0}v_r)\exp\left(-\ell\frac{k_{0e}k_{r0}v_e + k_{0r}k_{e0}v_r}{k_{0e}v_e v_r + k_{0r}v_e v_r}\right)}{v_e(v_r(k_{\text{on}}(k_{0e} + k_{0r} + k_{e0}) + k_{0r}k_{e0}) - v_e(k_{0e}(k_{\text{on}} - k_{r0}) + k_{\text{on}}(k_{0r} + k_{r0})))}, \tag{18}$$

$$\mathbf{P}_{\text{r}}(\ell) = \frac{k_{\text{on}}(k_{0e}k_{r0}v_e + k_{0r}k_{e0}v_r)\exp\left(-\ell\frac{k_{0e}k_{r0}v_e + k_{0r}k_{e0}v_r}{k_{0e}v_e v_r + k_{0r}v_e v_r}\right)}{v_e(k_{0e}(k_{\text{on}} - k_{r0}) + k_{\text{on}}(k_{0r} + k_{r0}))) - v_r(v_r(k_{\text{on}}(k_{0e} + k_{0r} + k_{e0}) + k_{0r}k_{e0})}. \tag{19}$$

$$\begin{aligned}\mathcal{P}(\ell) &= \mathbf{P}_0(\ell) + \mathbf{P}_{\text{r}}(\ell) + \mathbf{P}_{\text{e}}(\ell) \\ &= \frac{k_{\text{on}}(k_{r0}v_e - k_{e0}v_r)(k_{0e}k_{r0}v_e + k_{0r}k_{e0}v_r)\exp\left(-\ell\frac{k_{0e}k_{r0}v_e + k_{0r}k_{e0}v_r}{k_{0e}v_e v_r + k_{0r}v_e v_r}\right)}{v_e(k_{0e}(k_{\text{on}} - k_{r0}) + k_{\text{on}}(k_{0r} + k_{r0}))) - v_e v_r(k_{0e} + k_{0r})(v_r(k_{\text{on}}(k_{0e} + k_{0r} + k_{e0}) + k_{0r}k_{e0})}. \end{aligned} \tag{20}$$

### 3 Simulation of T4P-based migration

#### 3.1 Model description

Bacteria are idealized as point-like objects, endowed with a director and moving on a plane. T4P production sites are randomly placed on opposite sides of the bacterium, according to experimentally determined distributions of pilus locations along the cell body. Individual T4P are modelled as Hookean springs whose rest lengths are being extended or retracted according to the stochastic process depicted in Fig. 2a of the main text. The rest length of the $i^{\text{th}}$ pilus, $\ell_0^i$, changes over time as

$$\frac{d\ell_0^i}{dt} = \begin{cases} v_{\text{r}} < 0 & \text{if retracting,} \\ v_{\text{e}} > 0 & \text{if elongating,} \\ 0 & \text{if idle,} \end{cases} \tag{28}$$

where the retraction speed can be depend on the load on the pilus, see below. Pilus sites becomes active with rate $k_{\text{on}}$ and start elongating pili in the direction $\mathbf{e}_i$. These direction vectors are generated randomly by drawing from a measured distribution of angles between pili and the cellular long axis. The angular distributions shown in Fig 5d of the main text are aligned such that the pre-determined polarity of the simulated bacterium coincides with the dominant pole in the experimental data where pili genesis occurs most frequently. If, at any time, a pilus in the retraction state reaches a vanishing rest length, this pilus site becomes inactive until it stochastically switches on again. If the pilus is attached to the surface during a switch to inactivity, it is forced to detach. Any active pili that are have rest lengths $\ell_0 \geq 0.25\,\mu m$ can bind to the substrate with rate $\pi_p$. Attached pili detach from substrate with a force-dependent rate $\lambda_p$. When attached, the tension on the $i^{\text{th}}$ pilus ($i \in \mathcal{A}_{\text{p}}$) with rest length $\ell_0^i$ and length $\ell^i$ is $\mathbf{F}_i^{\text{p}} = \kappa_p \max\left[(\ell^i - \ell_0^i), 0\right]\mathbf{e}_i$ where $\kappa_p$ is the spring constant. Note that attached pili buckle if their rest length is greater than the distance between tip attachment position and bacterium position. Also, the retraction speed depends on the force on the pilus as $v_{\text{r}}(F_i^{\text{p}}) = v_0(1 - F_i^{\text{p}}/F_{\text{stall}})$ where $v_0 < 0$ is constant.

Passive adhesion bonds are modelled as Hookean springs with a vanishing rest length and they act as anchors, holding the bacterium to the surface and sometimes opposing the tension generated by the pili. Passive adhesins bind to the surface with rate $\pi_{\text{pa}}/C_{\text{red}}$, where the factor $C_{\text{red}}$ takes into account that the the cell body of twitching bacteria occasionally fully detaches from the substrate. Once all passive adhesion bonds are broken, the larger space between the cell and the substrate likely reduces the rate of formation of new passive adhesion bonds, which we model by setting $C_{\text{red}} = 1$ if any passive adhesion bond is already formed and $C_{\text{red}} \geq 1$ when all passive adhesion bonds are broken. Breakage of passive adhesion bonds occurs with a force-dependent rate $\lambda_{\text{pa}}$. The $i^{\text{th}}$ passive adhesion ($i \in \mathcal{A}_{\text{pa}}$) transmits a force $\mathbf{F}_i^{\text{pa}} = \kappa_a s_i^a \mathbf{e}_i^{pa}$ where $s_i^a$ is the stretch and $\mathbf{e}_i^{pa}$ is the orientation of the passive adhesion that is taken to be opposite to the cell's migration direction after first attachment. All direction vectors of attached pili and passive adhesions are updated at every simulation time step $\Delta t$ based on the displacement of the bacterium and the surface-attachment points of the pili.

Cell-wide force balance is calculated with an equation for overdamped motion, where the instantaneous velocity of bacterium $\dot{\mathbf{x}}(t)$ at time $t$ is determined by

$$\eta\,\dot{\mathbf{x}}(t) = \sum_{i\in\mathcal{A}_{\text{p}}} \mathbf{F}_i^{\text{p}} + \sum_{i\in\mathcal{A}_{\text{pa}}} \mathbf{F}_i^{\text{pa}}, \tag{29}$$

with $\eta$ being the friction coefficient. Detachment rates of T4P and passive adhesions are modeled as force-dependent slip bonds.

**Simulation of mutant cells** In order to simulate different mutants, the T4P creation rate $k_{\text{on}}$ is changed. ΔpilA cells are simulated by setting the rate for new pilus creation to zero, $k_{\text{on}} = 0.0\,\text{s}^{-1}$. Δ*pilH* mutants are simulated with

Table S2: Parameters for the simulations (orange lines in Figs. 2,3,5, and 6 of the main text). Parameters governing T4P statistics were matched to experimental data for bacteria on 55 kPa PAA substrates. Where a range of parameters is given, values were adjusted to match migration behavior on different substrates. The time scale is $\tau_0 = 1$ sec, the length scale is $L_0 = 1\ \mu$m, and the stiffness scale is $\kappa_0 = 2000\,$pN$/\mu$m based on [3].

| **Parameter** | **Symbol** | **Value** | **Notes** |
|---|---|---|---|
| Simulation time step | $\Delta t$ | $10^{-4}\ \tau_0$ | chosen based on biggest rate |
| Total simulation time steps | | $> 6 \cdot 10^6$ | including $10^6$ discarded initial steps |
| Pili spring coefficient | $\kappa_p$ | $1\ \kappa_0$ | |
| Pas. adh. spring coef. | $\kappa_a$ | $1\ \kappa_0$ | chosen as such for simplicity |
| Number of pilus sites | | 30 | Determines max # of pili. |
| Number of pas. adh. | $N_{pa}$ | [5-7] | |
| Pili retraction velocity | $v_r$ | $-0.55\ L_0/\tau_0$ | |
| Pili elongation velocity | $v_e$ | $0.42\ L_0/\tau_0$ | |
| Pili stall force | $F_{\text{stall}}$ | $0.05\ K_0 L_0$ | |
| Threshold for contribution | $F_{\text{thresh}}$ | $0.0025\ K_0 L_0$ | $\simeq$ 5pN |
| Friction coefficient | $\eta$ | $20\Delta t\ \kappa_0$ | This choice allows for a fast relaxation of visous forces. |
| Fraction of pili at the front | | 68% | genesis angle in $[-\pi/3, \pi/3]$ |
| Fraction of pili at the back | | 12% | genesis angle in $[2\pi/3, 4\pi/3]$ |
| Fraction of pili at the sides | | 20% | genesis angle in $[\pi/3, 2\pi/3] \cup [4\pi/3, 5\pi/3]$ |
| Pili binding rate | $\pi_p$ | $[0.5 - 2.3]\ \tau_0^{-1}$ | |
| Pas. adh. binding rate const. | $\pi_{\text{pa}}$ | $[1 - 2.3]\ \tau_0^{-1}$ | |
| Pili rupture rate const. | $k_r = \lambda_{\text{p}}(0)$ | $[0.75 - 1]\ \tau_0^{-1}$ | |
| Pas. adh. rupture rate const. | $k_r^{\text{pa}} = \lambda_{\text{pa}}(0)$ | $[0.75 - 1]\ \tau_0^{-1}$ | |
| Force sensitivity of T4P bond | $F_{\text{sens}}$ | $0.025\ K_0 L_0$ | $\simeq$ 50pN ($\lambda_{\text{p}}(F) = k_r \exp F/F_{\text{sens}}$). |
| Force sens. of pas. adh. bond | $F_{\text{sens}}^{\text{pa}}$ | $0.005\ K_0 L_0$ | $\simeq$ 10pN ($\lambda_{\text{pa}}(F) = k_r^{\text{pa}} \exp F/F_{\text{sens}}^{\text{pa}}$). |
| Pas. adh. binding reduction | $C_{\text{red}}$ | $[1 - 4.5]$ | use $C_{\text{red}} = 1$ any Pas. adh. bond already formed |
| *Pilus rates:* | | | |
| T4P site activation | $k_{\text{on}}$ | $[0.27 - 0.36]\ \tau_0^{-1}$ | based on the data displayed in Fig. S4h |
| Elongation to idle | $k_{\text{e0}}$ | $0.95\ \tau_0^{-1}$ | see the main text |
| Retraction to idle | $k_{\text{r0}}$ | $0.1099\ \tau_0^{-1}$ | from Ref. [2] |
| Idle to elongation | $k_{\text{0e}}$ | $0.4167\ \tau_0^{-1}$ | from Ref. [2] |
| Idle to retraction | $k_{\text{0r}}$ | $2.50\ \tau_0^{-1}$ | from Ref. [2] |

$k_{\text{on}} = 0.918\,\text{s}^{-1}$, which is obtained from measured pili production [2], and a reduced passive adhesion binding rate $\pi_{pa} = 0$ since the majority of these mutants is standing vertically on the substrate in experiments without shear flow. For simulation of $\Delta pilH$ mutants, we assume an almost iotropic distribution of pili around the cell with only 10% of pili predominantly pointing preferentially along one, arbitrarily chosen, direction. We also found that simulation of the high motile $\Delta pilH$ mutants required the addition of Gaussian noise white noise to Eq. (29). This noise is assumed to only play a role when cells are fully detached, i.e., when no pili are bound to the substrate, and we employ a diffusion coefficient of $0.217 \times 2/3\,\mu\text{m}^2/s$ for the two-dimensional random walk.

Simulations with shear flow To simulate the effect of fluid shear forces acting on the adherent bacteria, a constant force $\mathbf{F}_{\text{shear}}$ is added to Eq. 29. We assume that the hydrodynamic force points in the opposite direction as the pilated cell pole, consistent with the observation that *P. aeruginosa* tends to align its long axis with the flow [4]. The magnitude of $\mathbf{F}_{\text{shear}}$ is estimated as a product of the given shear stress and an effective area, see [5]. Since *P. aeruginosa* is approximately $0.55 \pm 0.05\,\mu$m in width and $[1.5, 2.1]\mu$m in length [6], the effective area has a range $[0.2 - 1.0]\,\mu\text{m}^2$. Due to the horizontal alignment of the long axis with the flow direction, we assume an effective area of $0.6\,\mu\text{m}^2$ in all simulations. Cell detachment under flow is a highly complicated stochastic process and a proper simulation of the involved hydrodynamics is beyond the scope of this work. To nevertheless analyze the detachment process qualitatively, we take the probability of cell detachment to be proportional to the mean time that passes in simulations until all pili and passive adhesins are detached from the surface. A proportionality factor is used for converting this mean time to the mean time until cell detachment $\langle\tau\rangle$ (320 for WT cells and 800 for $\Delta PilH$ cells).

#### 3.2 Simulation of rod-like cells

To confirm the simulation results obtained with an idealized model where the elongated cell shape is neglected, we also study a rigid-rod model. In this model, the translational and rotational friction coefficients depend on the aspect ratio of the particle. For motion parallel and normal to the long axis with length $L_r$ we employ $\eta_{||} = \eta L_r$ and $\eta_\perp = 2\eta_{||}$, respectively. The rotational friction coefficient is given by $\eta_\theta = \eta_{||} L_r^2/6$ [7, 8]. The angle between pili and the bacterium's long axis are distributed according to a predefined distribution and passive adhesion bonds are formed at the center of the particle. Measurement noise is not included in the simulations. Rod-like bacteria are simulated for long times to investigate the effect of reorientation on the MSD and to confirm that we can use dot-like model for the time scale of the measure data, see Fig. S3.

##### 3.3 Parameter values in simulations

In order to contribute to twitching, a pilus must attach to the substrate and retract. To determine the rate constants for substrate-attachment and subsequent T4P retraction, we scanned parameter values in simulations to obtain the measured ratio of contributing pili, see Fig. S9. The binding rate constant of pili to PAA and PEGDMA is found to be in the range of $[0.2, 0.5]\,\mathrm{s}^{-1}$. In the experiments, we observe that contributing pili usually detach from the substrate within a few frames. Therefore, the mean rupture time is estimated to be in the range of $[0.5, 2]\,\mathrm{s}$. Parameter values governing the passive adhesion dynamics are inferred indirectly from analysis of twitching migration, see below. For simulations, parameter values for twitchers on 55 kPa PAA substrates are used (see Tbl. S2).

##### 3.4 Are passive adhesions undetected T4P at the cell rear?

In principle, adhesion forces hindering motion could also result from undetected short T4P at the rear end of the bacteria. For excluding this possibility, we simulate bacteria without passive adhesions. Although the simulated T4P dynamics faithfully represent the experimental observations, the MSDs calculated from simulations do not match the experimental results without the incorporation of passive adhesions, see Fig. S8.

##### 3.5 Analysis of simulation results

To compare pilus lengths in simulations and experiments, we coarse-grain the pilus length $\ell$ measured in simulations. Note that $\ell$ is the actual length, not the rest length as that is not observable. A coarse time scale of $t_{cg} = 0.5\,\mathrm{sec}$ is chosen since our experimental data is recorded at two frames per second. Temporal coarse graining of the $i^{\mathrm{th}}$ pilus produces lengths as

$$\tilde{L}^i_{cg}(t) = \frac{\Delta t}{t_{cg}} \sum_{t}^{t+t_{cg}} \ell(t) \ . \tag{30}$$

To emulate the measurement error resulting from the finite spatial resolution of the microscopy images, Gaussian random noise with vanishing mean and a standard deviation of 0.05 is added to $\tilde{L}^i_{cg}(t)$ where $\tilde{L}^i_{cg} > 0$, thus excluding non-visible pili. Extension and retraction speeds are calculated from the coarse-grained data as $v^i_{m_{cg}}(t_m) = \frac{L^i_{cg}(t+t_{cg}) - L^i_{cg}(t)}{t_{cg}}$ .

MSDs and ensemble-averaged MSDs are calculated from the simulation data the same way as for experimental data, where we sample the bacterial position every $(2t_{cg})$ since microscopy images of twitching cells are recorded at one frame per second. Measurement errors in both components of the two-dimensional position vector are emulated by addition of random values drawn from an exponential distribution with mean $0.03\,\mu$m. Twitching velocity is calculated as $\mathbf{v}_{cg}(t) = (\mathbf{r}\,(t + (2t_{cg})) - \mathbf{r}\,(t))\ /\ (2t_{cg})$, where $\mathbf{r}(t)$ is the position of the bacterium at time $t$.

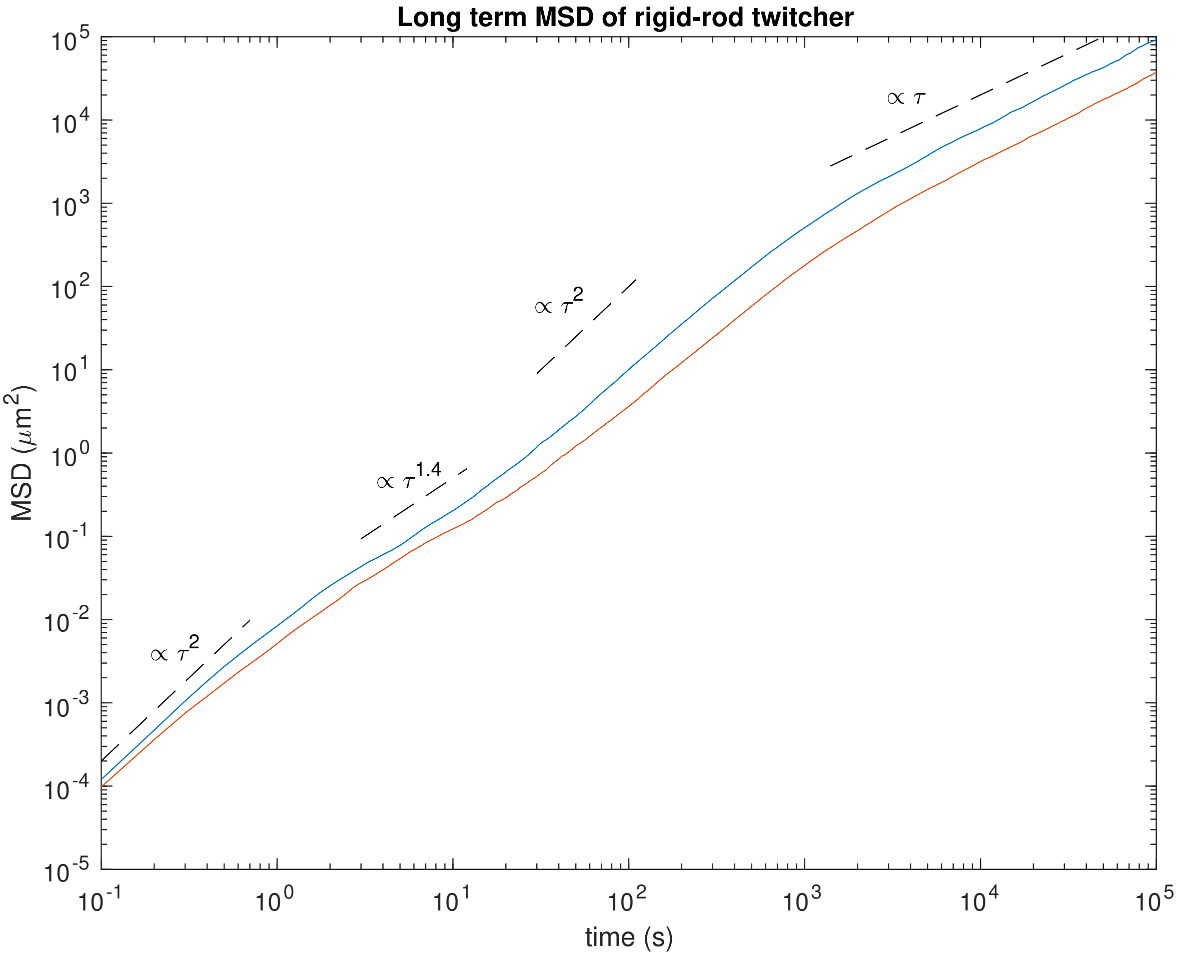


Figure S3: Simulated MSD for unipolar bacteria, as in Ref. [9], except the pili appear in a 120 degrees cone instead of 90 and cells with the measured pili placement in this work are compared. Very long term, $>$ 1000 sec, MSD $\langle x - x_0 \rangle_{\text{ensemble}}$ of rigid-rod twitchers with direction as a degree of freedom and without passive adhesions, see Methods. 1000 simulations per line.

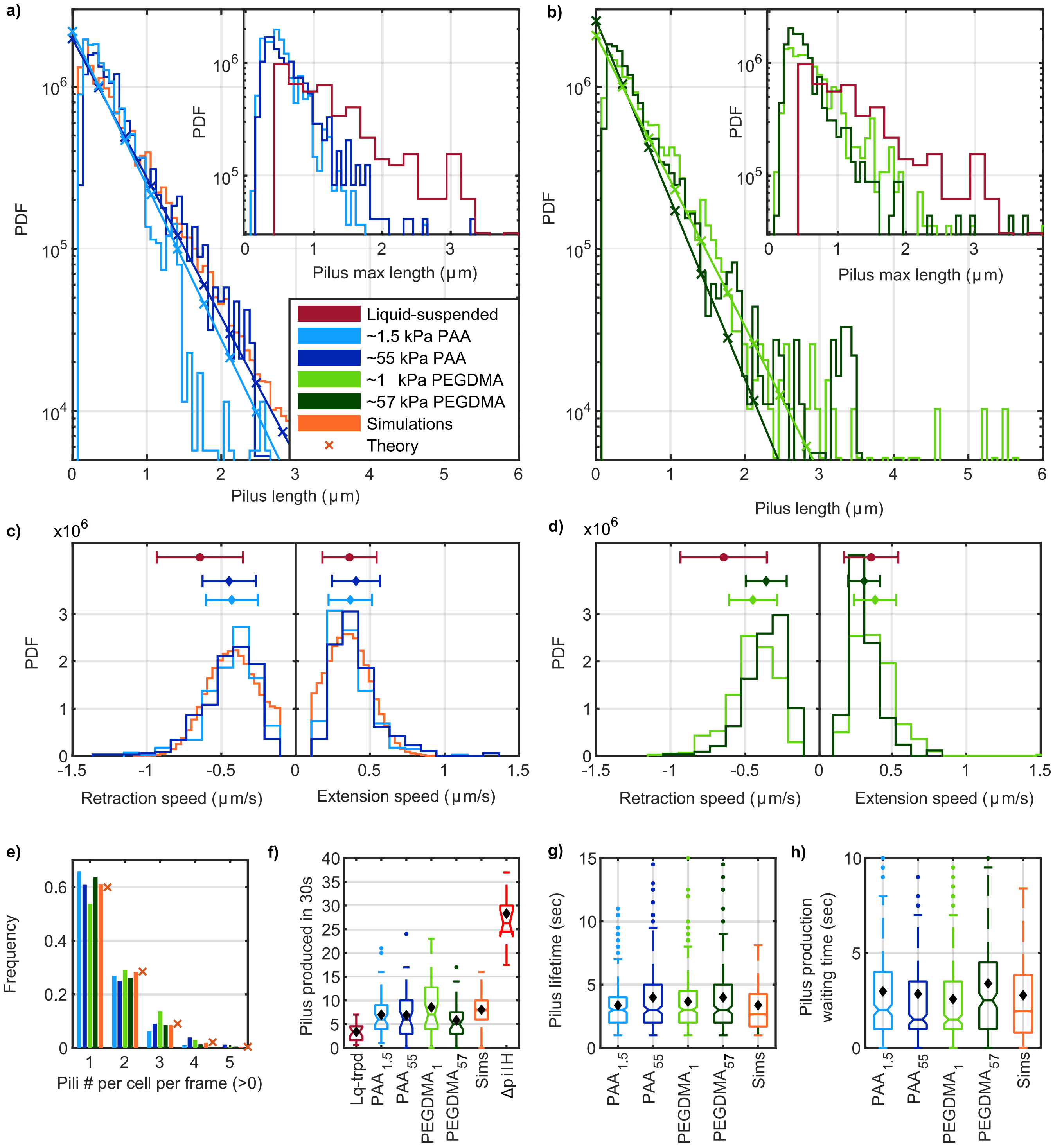


Figure S4: T4P dynamics are quantitatively explained by a stochastic 4-state model. Dynamics depend only weakly on the properties of hydrogel substrates, but pilus production is upregulated on surface contact. a-b) T4P length distributions for different substrates. Insets: distributions of maximum length during full extension-retraction cycles. Theory predicts exponential distributions as $\propto e^{-K\ell}$, with $K$ in Eq. (10), that differ only by the motor-speed parameters, which are taken from the means of the distributions in c-d. c-d) Distribution of mean retraction and extension speeds per pilus. Error bars show the mean and the standard deviation. e) Number of pili per cell per frame. Zeros are excluded since only cells with pili are considered. Crosses show the theoretically predicted, zero-truncated Poisson distribution, Eq.(27). f-h) Box plots of T4P statistics, black diamonds mark the mean of the data. f) Number of T4P produced in 30 seconds. Lower T4P production for cell in liquid suspension is significant ($p = 3.5 \times 10^{-9}$, Welch's t-test). g) Lifetime of T4P between first appearance and full retraction, h) time between appearance of new T4P. Data for liquid-suspended cells (dark red) is from [2]. Simulation are performed with parameters corresponding to T4P statistics on 55 kPa PAA gels. See appendix. To emulate experimental constraints (average pilus lifetime is expected to be less, see pilus lifetime in methods), short-lived pili ($< 1$ s) are omitted from simulation data. See text regarding details of the box plots.

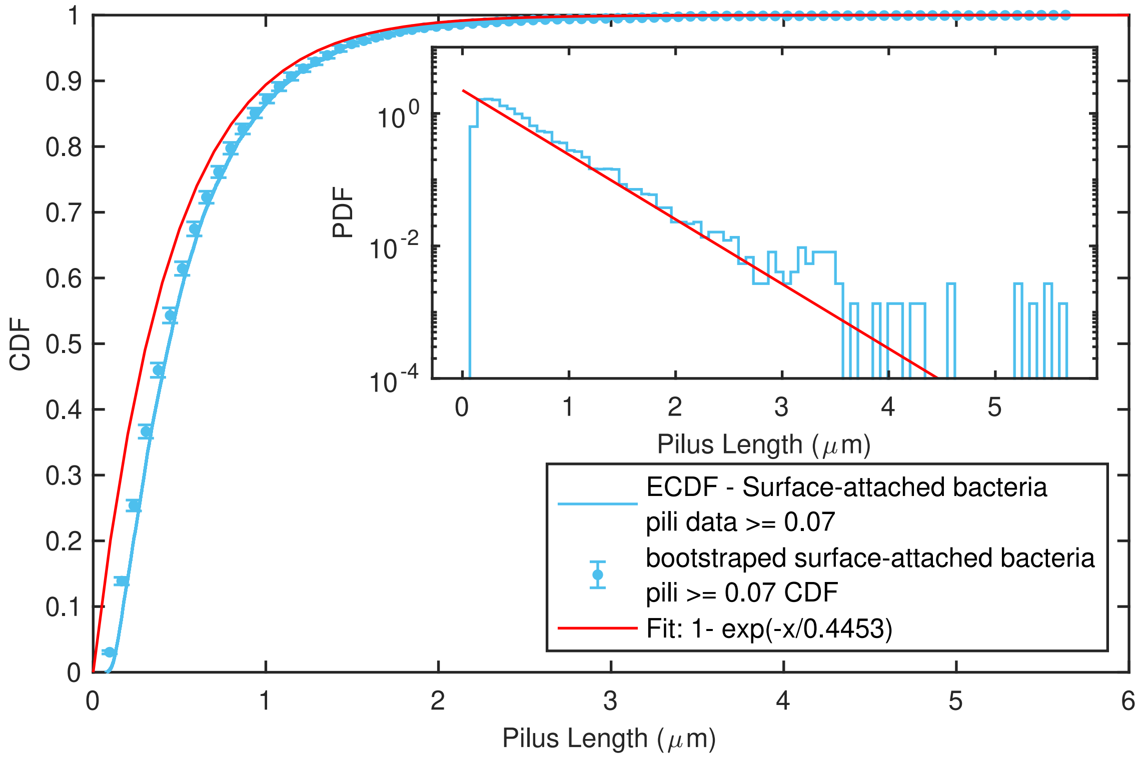


Figure S5: Using MATLAB's built-in curve-fitting tool (cftool), a custom equation $1 - \exp(-x/\mu)$ is fitted to bootstrapped pilus length CDF (10000 times). The mean is found as $\mu = 0.4453$ with 95% confidence bounds (0.4443, 0.4464), and SSE is 0.0001808. ($0.07\mu$m = 1pixel).

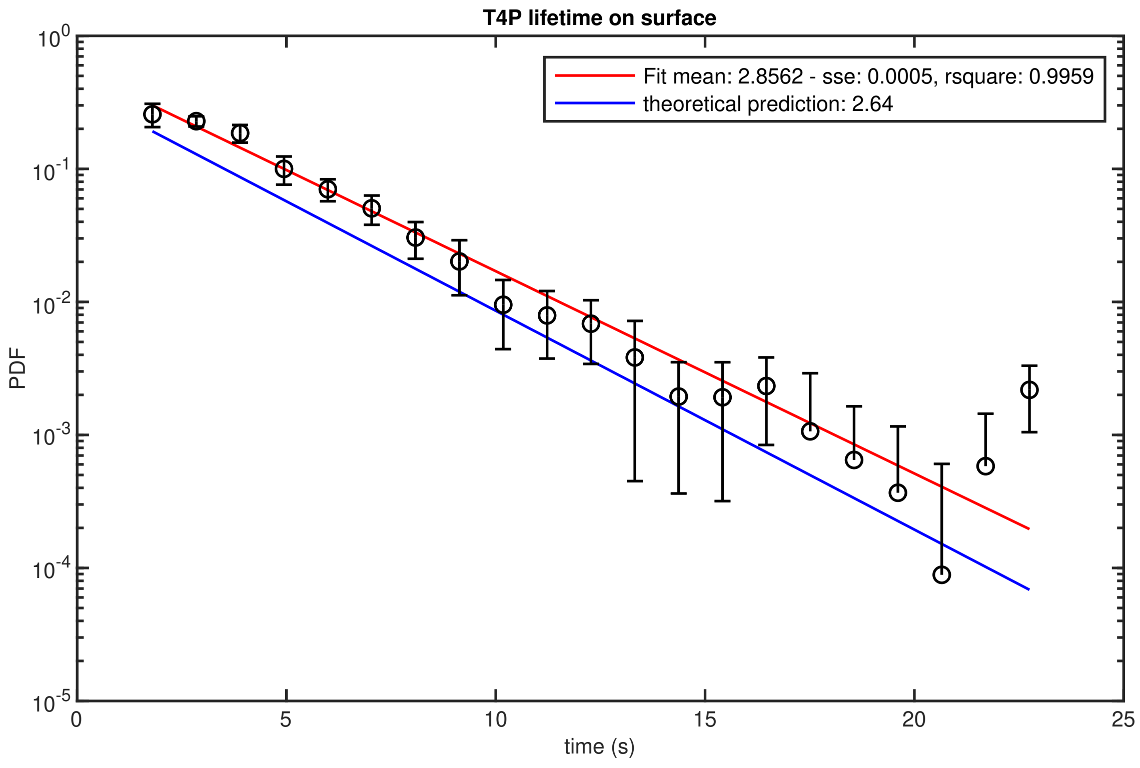


Figure S6: The fit ($c \exp(-x/\mu)$) is calculated automatically by MATLAB using non-linear least squares (with robustness LAR) to bootstrapped PDFs of pilus lifetime data.

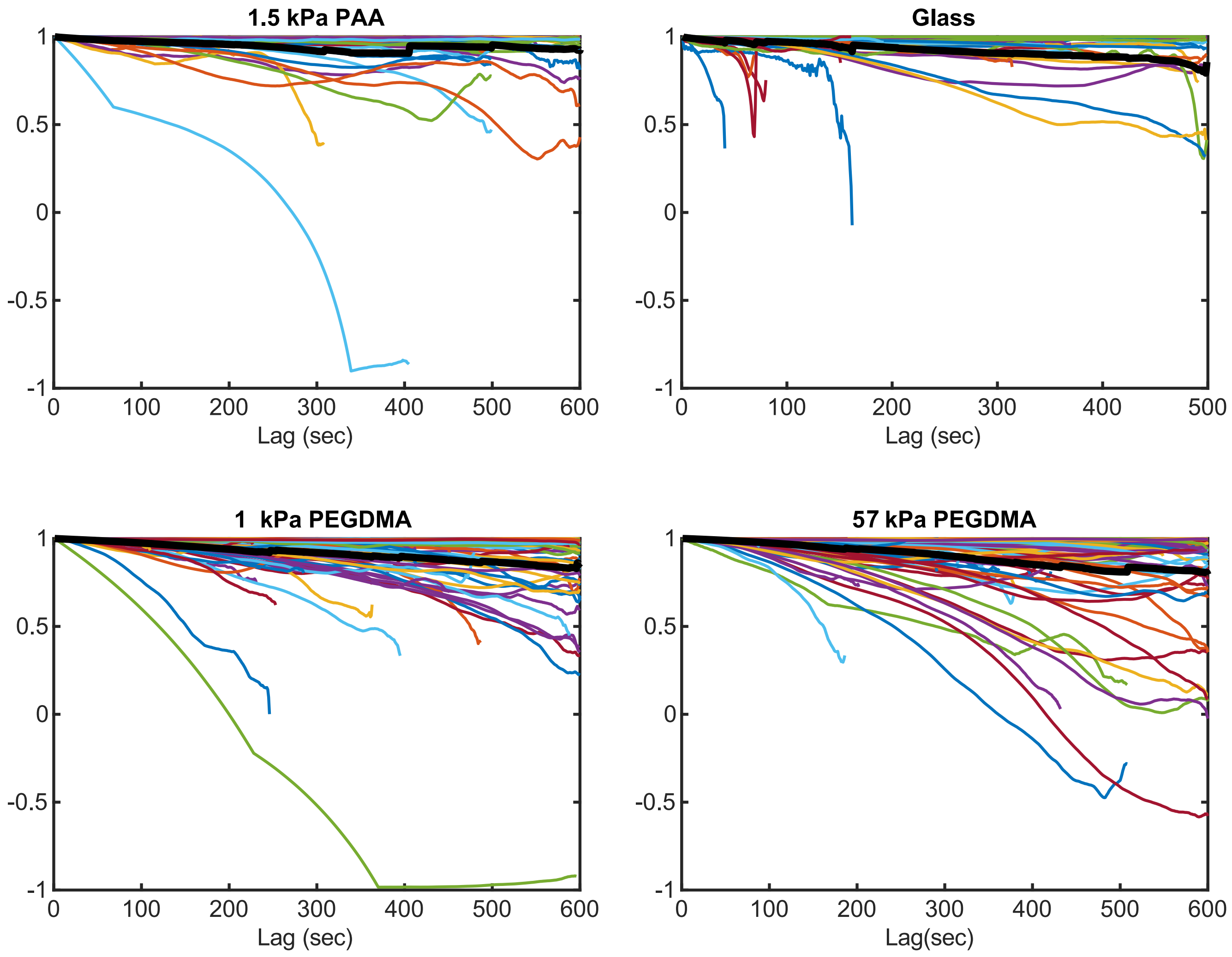


Figure S7: Autocorrelation of the orientation the long axis of tracked bacteria.

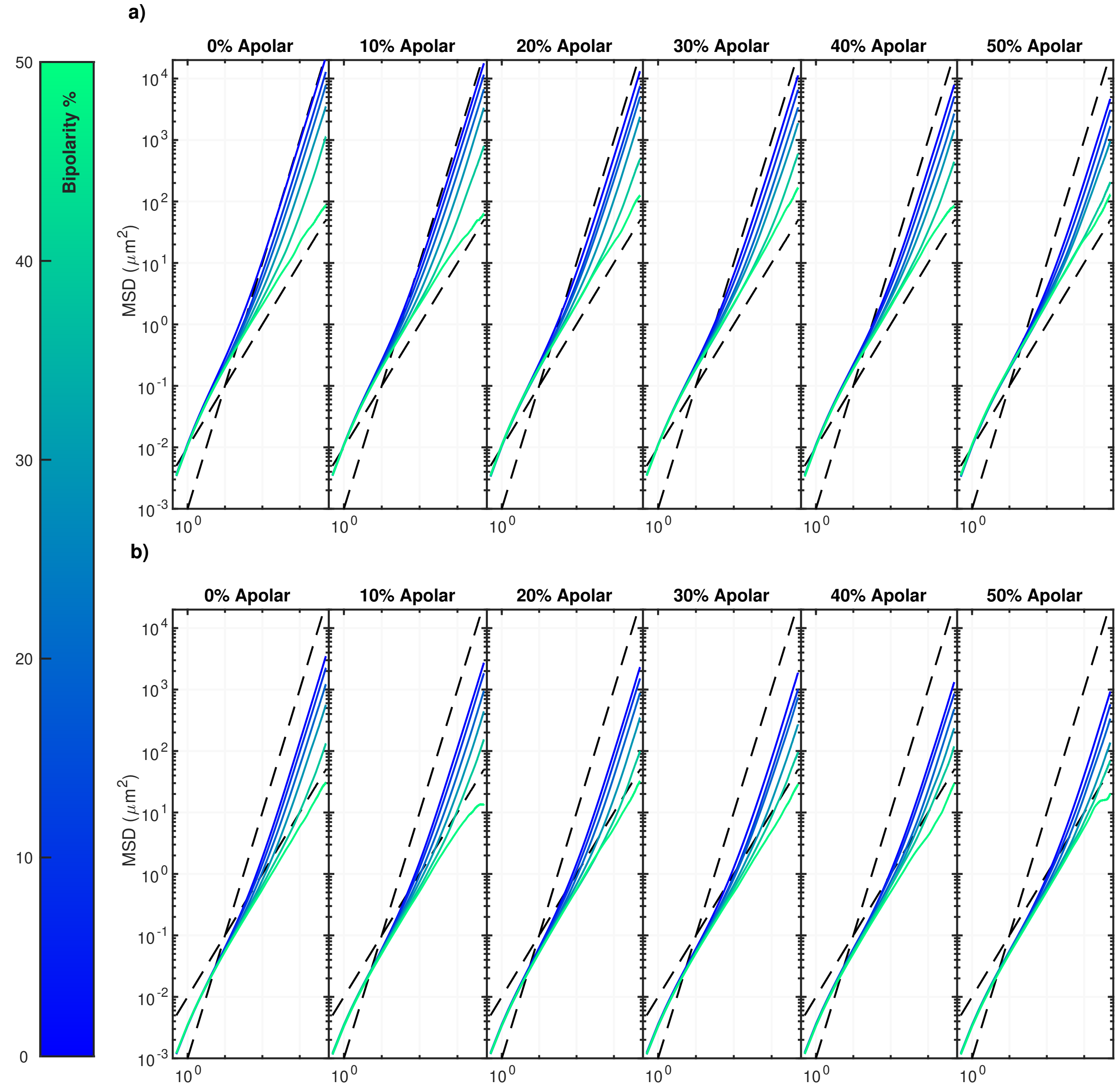


Figure S8: Pili distribution and migration dynamics. Bipolarity and apolarity impacts the duration of the crossover regime. Bipolarity also impacts long time behaviour. Simulations in this figure are performed with parameter values measured for 55 kPa PAA except the cells do not have any passive adhesions. (5 simulation per line.) Panel a with ≈ 30% and panel b with ≈ 20% contributing pili.

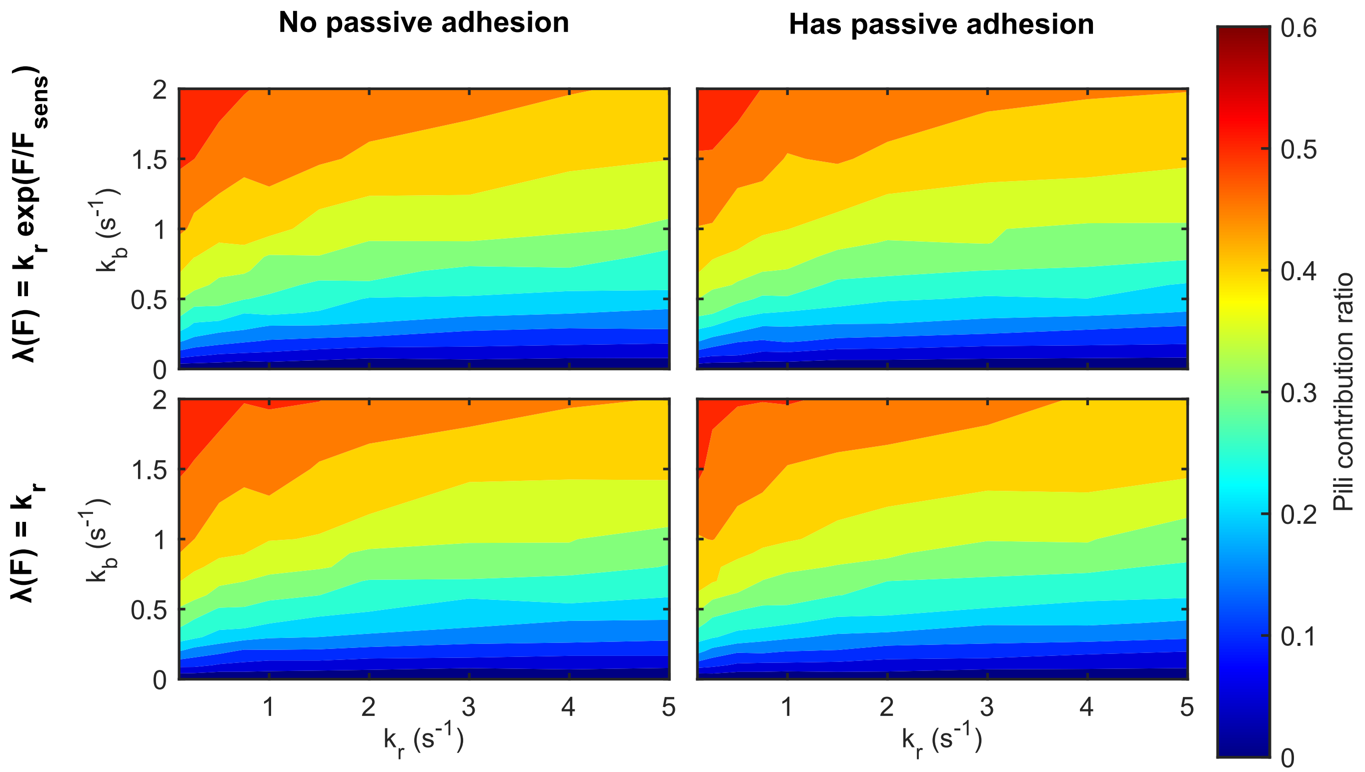


Figure S9: State diagram showing contributing pili ratios for varying binding and rupture rates between T4P and the substrate.

Table S3: Number of measured samples by figure. All data were collected from at least three experiments. Numbers follow the scheme: $(\mathrm{PAA}_{1.5}, \mathrm{PAA}_{55}, \mathrm{PEG}_{1}, \mathrm{PEG}_{57}, x, y)$ where x and y are described in notes.

| Fig. | Panel | # of bacteria | # of samples | Notes |
|---|---|---|---|---|
| S4 and | a, b | (49,41,39,45) | (3194,3313,3481,3147) | Dead pili are not included. |
| 2 as its subset | | | (2505,2716,2773,2597) | Data bigger than 1 pixel |
| | a&b insets | (49,41,39,45, ) | (242,228,398,320,307) | x: Lq. trpd. |
| | c, d (S14) | (49,41,39,45, -) | (387,343,398,320, -) | x: Lq. trpd. [2] |
| | e, f | (49,41,39,45,24,10) | (49,41,39,45,24,10) | x: Lq. trpd., y: ΔPilH |
| | g | (49,41,39,45) | (304,255,311,236) | |
| | h (S1) | (49,41,39,45) | (294,244,296,219) | |
| 3 | b | (28,31,25,37) | (50,98,49,81) | |
| | c | (49(28),41(31),39(25),45(37)) | (50,98,49,81) | inner: the bacteria that had contributing pili |
| 5 | a, b, e | (55,67,101,102, 69) | | x: ΔpilH |
| | c | 67 | | |
| | d | (49,41,39,45) | | |
| 6 | WT | | (500, 537, 383, 266, 100) | (1, 4, 14, 37, 123 ) kPa |
| detached | ΔPilA | | (63,137,340,369,604) | (1, 4, 14, 37, 123 ) kPa |
| number of cells | ΔPilH | | (462, 349, 131, 267, NA) | (1, 4, 14, 37, NA ) kPa |
| S2 | | (25,21,23,24) | | |
| S5 | | 10591 | | |
| S6 | | 1106 | | |
| S7 | | (55,- ,101,102, 45) | (55,- ,101,102, 45) | x: Glass |
| S10 | top | (338-49,255-88,341-57,231-89) | | NC-C |
| | bottom | (265-39,193-62,269-42,177-59) | | NC-C |

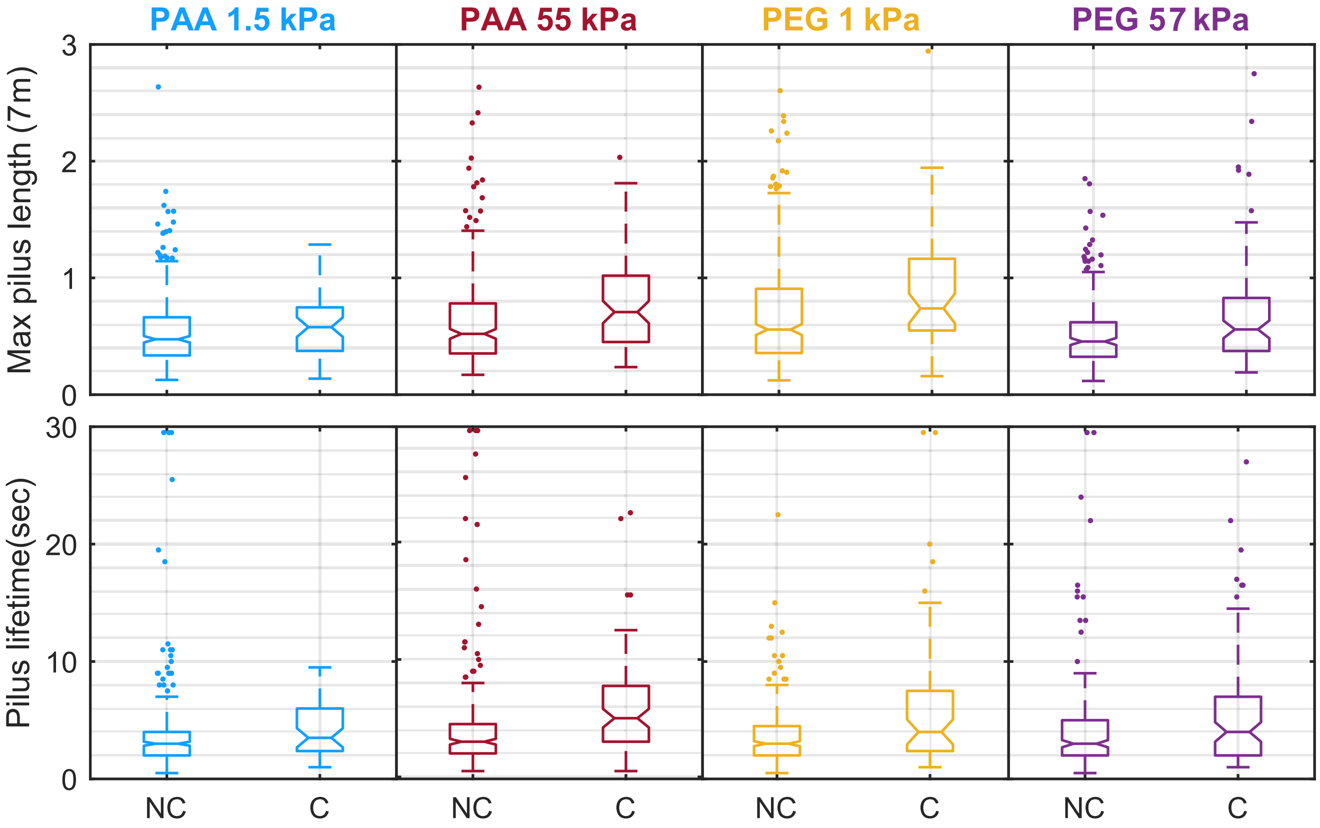


Figure S10: Pili with long length and lifetime have a high chance of adhering to surfaces. Non-contributing pili spend more time for extension than for retraction on all gels.

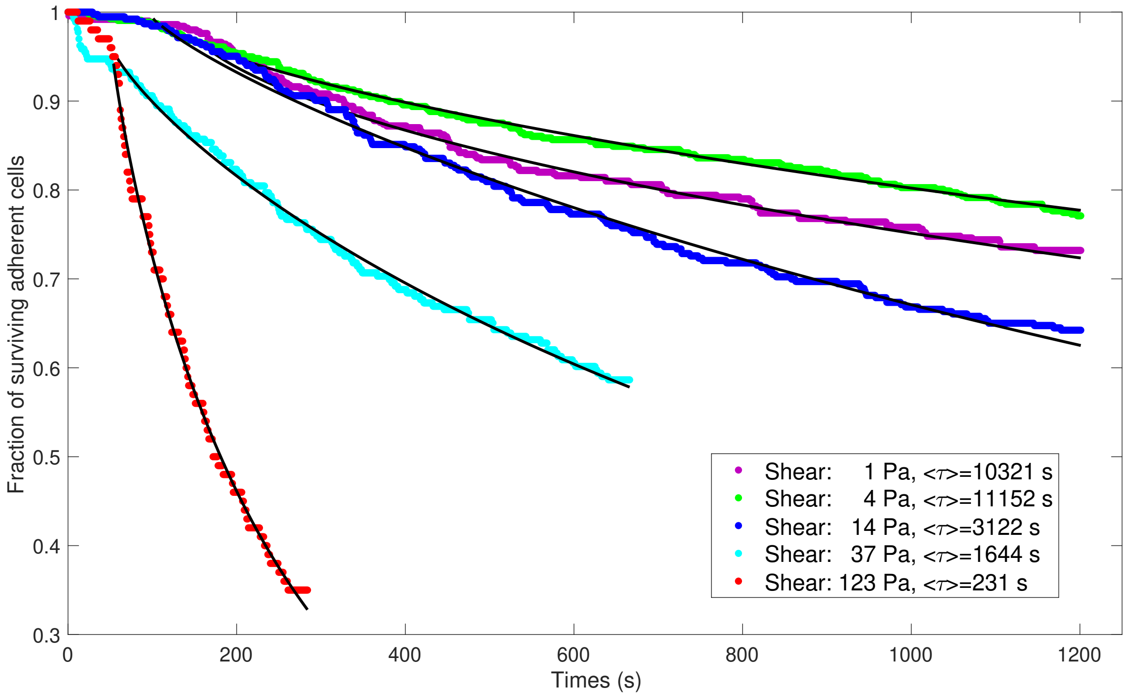


Figure S11: Fractions of WT cells that remain attached for a given time after onset of the shear flow. The fraction of cells surviving attached to the surface is fitted with a stretched exponential function $a\exp[-((t-t_0)./\tau)^{\alpha}]$. Here, $t_0$ is used to incorporate a time lag between turning on the syringe pump and the establishment of stationary flow during wich cells experience various levels of shear ($t_0$ is around 100s ). $\tau$ represents a timescale of survival and $\alpha \in [0,1]$ is the stretch exponent. All distributions were created from three separate experiments, each measuring detachment of statistics for 100 to 1000 cells.

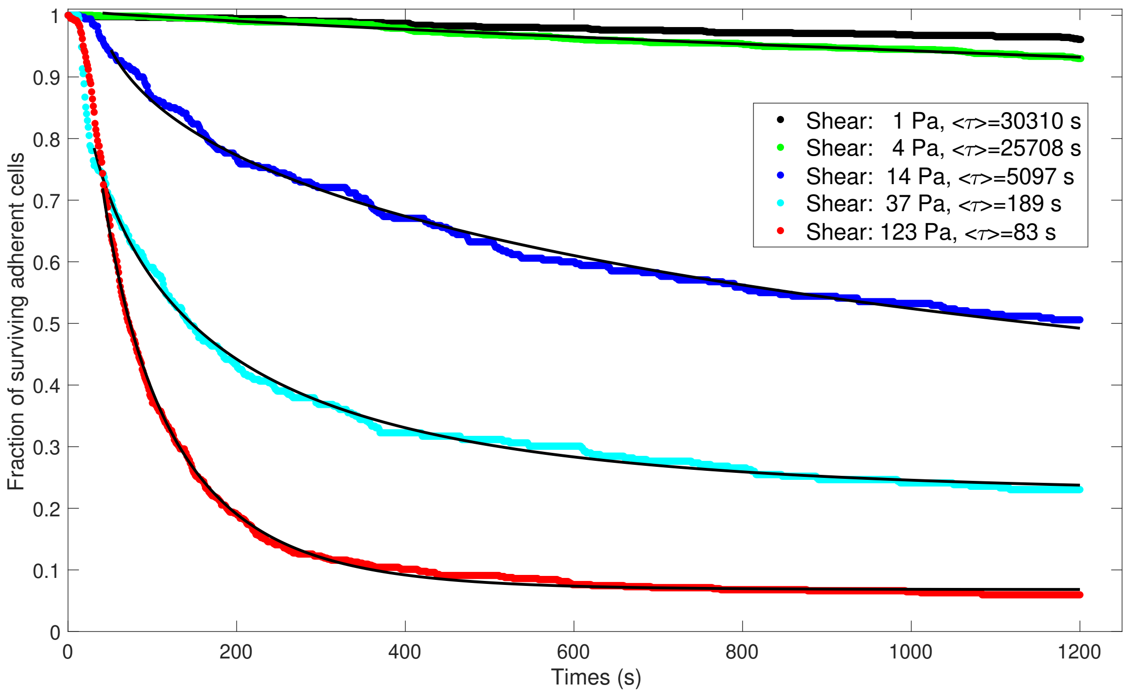


Figure S12: Fractions of *pilA* mutants that remain on the surface for a given time after onset of the shear flow. These mutants are devoid of T4P. For shear magnitudes $\geq 14\,\mathrm{Pa}$, we observe for these mutants a pronounced downstream sliding motion with a slip-stick character prior to detachment. In this data, cells are only considered detached if they leave the surface completely. Like for WT cells, the fraction of mutants that survives on the surface is fitted with a stretched exponential function. All distributions were created from three separate experiments, each measuring detachment of statistics for 100 to 1000 cells.

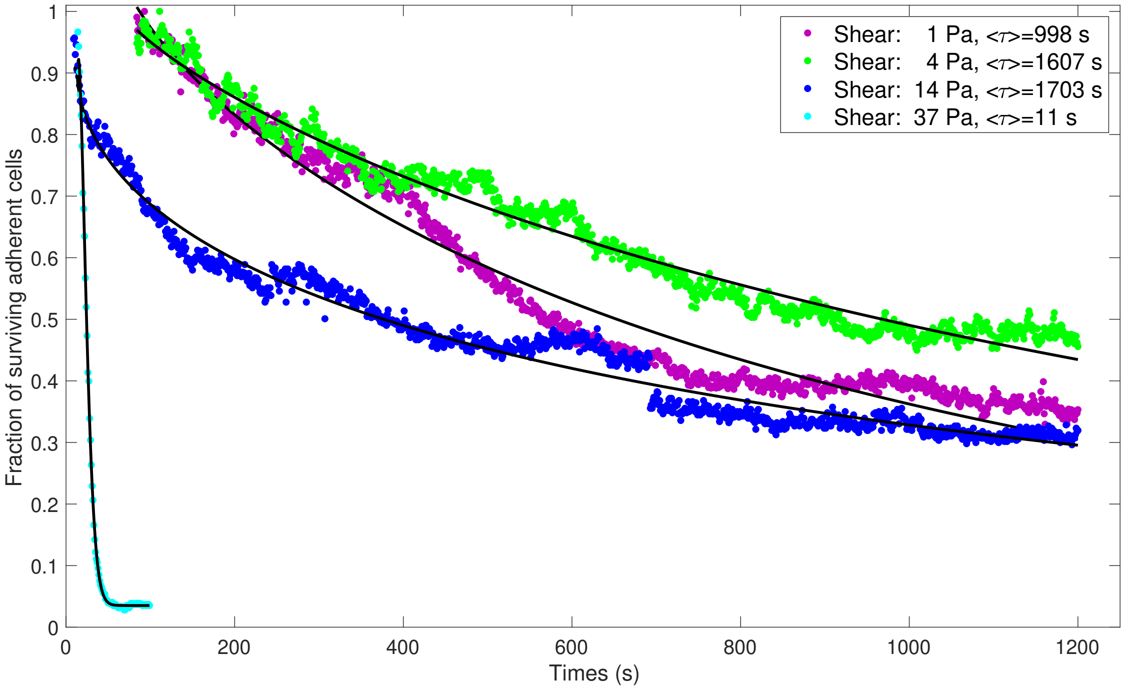


Figure S13: Fractions of *pilH* mutants that remain on the surface for a given time after onset of the shear flow. These mutants generate an exessive number of T4P and preferentially attach with one pole to twitch in an upright orientation. For shear magnitudes $\geq$ 14 Pa, we observe for these mutants a pronounced downstream sliding motion. In this data, cells are only considered detached if they leave the surface completely. Like for WT cells, the fraction of mutants that survives on the surface is fitted with a stretched exponential function. All distributions were created from three separate experiments, each measuring detachment of statistics for 100 to 1000 cells.

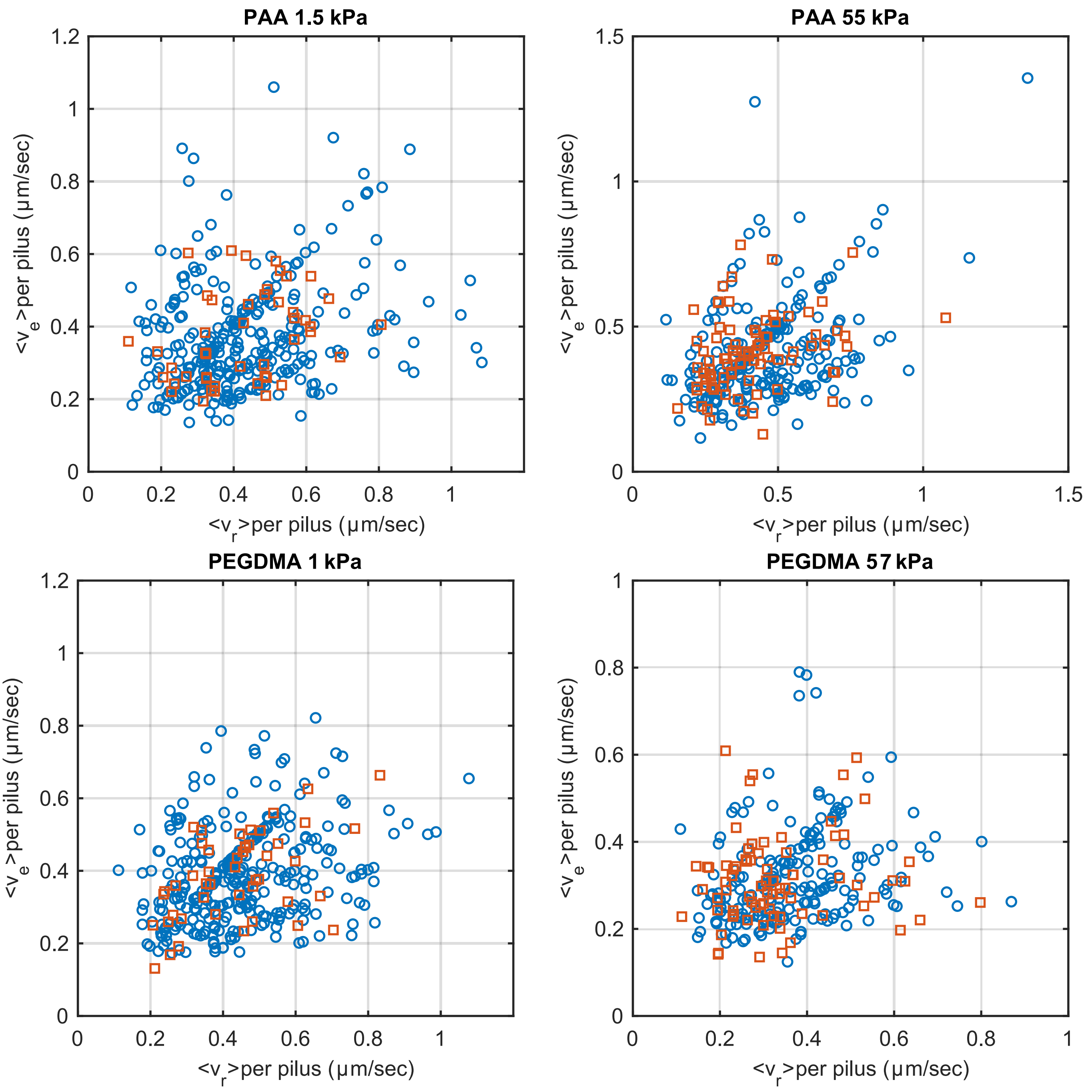


Figure S14: The average retraction speed against the average elongation speed per pilus. Squares represent contributing pili and circles represent the not contributing pili.
